# Genome-wide association study of frost tolerance in *Vicia faba* reveals syntenic loci in cool-season legumes and highlights relevant candidate genes

**DOI:** 10.1101/2024.11.27.624268

**Authors:** Baptiste Imbert, Jonathan Kreplak, Isabelle Lejeune-Hénaut, Jean-Bernard Magnin-Robert, Gilles Boutet, Pascal Marget, Grégoire Aubert, Judith Burstin, Nadim Tayeh

## Abstract

Cool-season grain legumes are mostly grown over spring and summer due to poor frost tolerance. However, fall-sown varieties often provide higher yields, earlier harvests and avoid late-season drought and heat. Understanding the genetic determinism and molecular basis of frost tolerance is therefore crucial for developing high-performing winter varieties. This study aimed to (1) investigate the genetic architecture of frost tolerance in *Vicia faba* L. using 247 accessions phenotyped under four field environments, and (2) explore the conservation of frost tolerance loci in cool-season legumes using the OrthoLegKB translational research database. A genome-wide association study identified nineteen *V. faba* genomic regions with a high density of markers significantly associated with frost tolerance, on all chromosomes. Mapping of frost tolerance QTL from *V. faba* and related species obtained from the literature onto their respective reference genomes and their integration into OrthoLegKB revealed synteny of major QTL across *V. faba*, *Pisum sativum*, and/or *Medicago truncatula*, particularly near clusters of *CBF/DREB1* genes. Frost tolerance QTL at the *P. sativum Le* locus, which controls internode length, were also syntenic with a frost tolerance QTL in *V. faba*. Synteny between frost tolerance QTL and those controlling phenology and physiology was found at other loci, suggesting pleiotropy. Finally, expression data from *P. sativum* and *C. arietinum* accessions grown under low temperature were considered as information source to highlight potential candidate genes underlying the conserved QTL. Overall, these results provide a valuable resource for understanding and improving frost tolerance in *V. faba* and other cool-season legumes, including orphan crops by knowledge transfer. The use of OrthoLegKB to explore the genetic and molecular determinism of target traits across species is worth generalising.

## Introduction

Cool-season grain legumes are key components of sustainable cropping systems. These crops are valuable food and feed ingredients due to the high protein content and the health benefits of their seeds, such as reduced risk of cardiovascular and chronic diseases (Zhao et al., 2024). However, their development depends on their ability to withstand the many challenges they face in the dedicated agroecosystems, including abiotic and biotic stresses that can significantly affect yield. Faba bean (*Vicia faba* L.) and pea (*Pisum sativum* L.) are among the most yielding grain legumes worldwide (FAO, 2022), and their yield potential could be enhanced by increasing the proportion of cultivated winter varieties over spring varieties. Winter varieties are sown in autumn (Bourion et al., 2003; Link et al., 2010; Flores et al., 2013), allowing for a longer growing cycle that can optimise plant development and maximise yield potential (Klein et al., 2014). Despite a recent increase in the area dedicated to winter varieties, spring varieties remain the preferred option due to insufficient winter-hardiness in some agroclimatic conditions (Arbaoui et al., 2008; Lecomte et al., 2023). The process of cold acclimation, induced after exposure to low but not freezing temperatures, allows plants to survive the winter with minimal damage. Cold acclimation involves a number of changes in cell composition and structure that enable plants to resist subsequent frost episodes (Thomashow, 1999; Bourion et al., 2003). This includes changes in cell wall lipid composition and biosynthesis of cryoprotectants (sugars, amino-acids, etc.) that prevent the formation and expansion of ice crystals and help to withstand the mechanical stress exerted on the membranes (Ambroise et al., 2020). The expression of *C-repeat binding factor* (*CBF*)/*dehydration-responsive element binding factor 1* (*DREB1*) genes has been shown to be induced by low temperature and to be critical for frost tolerance in *Arabidopsis thaliana* (Thomashow, 1999) and in several crops (Liu et al., 2015; Würschum et al., 2017). *CBF/DREB1* genes encode for Apetala 2 (AP2)/ethylene responsive factor (ERF) transcription factors that promote the expression of cold-regulated (*COR*) genes and contribute to adapting the plant metabolism to survive cold. The *CBF/DREB1* signalling pathway plays a key role in frost tolerance, in parallel of a *CBF/DREB1*-independent pathways (Larran et al., 2023).

The intrinsic complexity of the control of frost tolerance *per se*, coupled with the prevalence of biotic stresses affecting plants under field conditions, has made the genetic analysis of frost tolerance in cool-season legumes a challenging endeavour. In *P. sativum*, several genomic regions regulating frost tolerance have been identified by quantitative trait loci (QTL) analyses using either diversity panels (Beji et al., 2020) or biparental populations (Klein et al., 2014), in particular the population derived from a cross between the frost-tolerant forage line Champagne and the frost-susceptible spring line Térèse (Lejeune-Hénaut et al., 2008; Dumont et al., 2009; Boutet et al., 2023). In the case of *V. faba*, the old French population Côte d’Or (Arbaoui et al., 2008; Sallam et al., 2016b) and the variety Hiverna (Carrillo-Perdomo et al., 2022) were used as frost tolerant parents to construct biparental populations and identify the genomic regions involved in the control of this trait. Genome-wide association studies (GWAS) have also been conducted using the Göttingen Winter Bean Population (GWBP), which was constructed from eleven founder lines to introduce genetic diversity (Ali et al., 2016; Sallam et al., 2016a, 2022). However, these studies have been largely hampered by the lack of marker density in genetic maps due to the absence of a reference genome sequence and to the size and complexity of the *V. faba* genome. Studies aiming at highlighting frost tolerance QTL have been scarcer in *Lens culinaris* (Kahraman et al., 2004, 2010), *Cicer arietinum* (Mugabe et al., 2019) and *M. truncatula* (Avia et al., 2013).

Given the close phylogenetic relationship and high synteny among grain legumes and with the model legume *M. truncatula*, translational research represents a great opportunity for boosting discoveries and breeding in this group of species. Previous studies have pinpointed syntenic frost tolerance QTL. The frost tolerance QTL on *M. truncatula* chromosome 6 including a cluster of 12 *CBF/DREB1* genes in its confidence interval (Tayeh et al., 2013b) has been shown to be syntenic to frost tolerance QTL on linkage group (LG) VI of *P. sativum* (Tayeh et al., 2013a; Klein et al., 2014; Beji et al., 2020) and LGI of *V. faba* (Carrillo-Perdomo et al., 2022). In order to facilitate the identification of syntenic QTL, and to determine the identity and expression profiles of the underlying candidate genes, we have recently developed the Ortho_KB framework with a dedicated instance for legumes called OrthoLegKB. The framework can process properly formatted data from the literature to extract and aggregate information into a knowledge graph (Imbert et al., 2023). OrthoLegKB includes the recently published long read-based genome assemblies of *V. faba* accession Hedin/2 (Jayakodi et al., 2023), *P. sativum* cv. ‘Caméor’ (‘Caméor’ v2 in prep.; for v1, see Kreplak et al., 2019), *L. culinaris* cv. CDC Redberry (Ramsay et al., 2021), *C. arietinum* CDC Frontier (Garg et al., 2022), and *M. truncatula* accession A17 (Pecrix et al., 2018). The database also includes phenological and morphological QTL from the literature (Imbert et al., 2023) for which the context sequences of flanking markers were successfully aligned onto the respective genome assemblies. Once integrated in the graph, all entities are represented by nodes, and gene nodes are linked to QTL nodes if they are included in their confidence interval. For GWAS outputs, only the closest gene to the peak marker is linked. Recent developments have been made in Ortho_KB for comparative genomics of syntenic QTL. Mainly, QTL annotation has evolved to take into account Planteome ontologies (Cooper et al., 2024), similarly to what was performed with RNA-seq data (Imbert et al., 2023). OrthoLegKB now supports the simultaneous analysis of QTL of different variables but related to the same trait; for example, connecting frost damage and winter survival to frost tolerance.

The aim of this study was to decipher the genetic architecture of frost tolerance in cool-season legumes, in particular pulses, in order to provide tools for their genetic improvement. A *V. faba* panel of 247 accessions, phenotyped under four winter field environments and genotyped at high density using exome capture technology, was used to perform GWAS for frost tolerance. The identified QTL were compared with QTL reported in *V. faba*, *P. sativum* and *M. truncatula* using OrthoLegKB. This approach, enabled by the synteny and orthology-based modules of this database was combined with the detection of genes from *P. sativum* and *C. arietinum* differentially expressed in frost-tolerant and susceptible lines. Altogether, the results revealed large cross-species conservation of frost tolerance-controlling loci and robust candidate genes. The use of OrthoLegKB facilitated multi-trait, multi-species queries, allowing the exploration of common control of frost tolerance, plant morphology and phenology across species.

## Materials and Methods

### Plant material and field phenotyping

A core collection of 247 *V. faba* accessions was selected from a large genetic resource collection described by Carrillo-Perdomo et al. (2019). This set of accessions represents the morphological and geographical diversity of *V. faba*. It consists of 88 cultivars or breeding lines and 159 landraces including major, minor and equina types. Leaves were collected from one plant per accession, flash frozen in liquid nitrogen and stored at −80°C before processing to DNA extraction. The tissues were then ground in liquid nitrogen using a pestle and mortar. Genomic DNA extraction was performed using Nucleospin PlantII minikit (Macherey-Nagel, Hoerdt, France) according to the manufacturer’s instructions. To assess frost tolerance of the panel, four different field trials were carried out, with a two-block design. All trials were conducted in France, in 2013-2014 (Bretenière, Bourgogne-Franche-Comté and Theix, Auvergne-Rhône-Alpes) and 2016-2017 (Bretenière, Bourgogne-Franche-Comté and Orsonville, Île-de-France). One row of 20 plants was sown for each accession in each experimental block. Frost damage was evaluated on a 0 to 5 scale after each frost episode as described by Carrillo-Perdomo et al. (2022). The survival rate of each accession was assessed as the percentage of emerged plants that had survived at the end of the winter season.

### Genotyping and SNP calling

SNP discovery and genotyping were performed using an exome capture protocol followed by sequencing as described in Aubert et al. (2023) for *P. sativum*. Briefly, normalised DNA samples were fragmented using Adaptive Focused Acoustics^®^ Technology (Covaris Inc., Massachusetts, USA) with a target size of 250 bp. DNA fragments were then subjected to a next-generation sequencing library preparation procedure consisting of end repair and Illumina adaptor ligation using the KAPA HTP kit (Roche, Basel, Switzerland). Individual index sequences were added to each library. The *V. faba* cv. Hiverna transcriptome assembly was used as a Refset for probe design. Exome capture design was performed based on this Refset, after removing redundancy, chloroplastic and mitochondrial sequences, and masking repeats. Sequences were captured using the Roche SeqCap EZ Developer Kit according to the manufacturer’s instructions. The efficiency of the sequence capture reaction was assessed by measuring the relative enrichment and loss of targeted and non-targeted regions before and after the sequence capture reaction, using qPCR. The captured samples were sequenced on the HiSeq4000 sequencing platform (Illumina, California, USA) using a paired end sequencing strategy of 2 reads of 150 bases each (approximately 10 million reads per accession). Sequenced reads were trimmed for adaptor sequence using cutadapt v1.8.3 (Martin, 2011). The GATK variant calling best practices pipeline was run on the Hedin/2 v1 genome assembly (Jayakodi et al., 2023) using elprep v5.1.3 (Herzeel et al., 2021), and GATK v4.2.6.1 (Van der Auwera and O’Connor, 2020) was used to merge SNP calling data and produce a final VCF file.

### SNP filtering and genome-wide association analysis

SNPs were first filtered for a heterozygosity rate of less than 15% using a custom script and bcftools (Danecek et al., 2021). Then, SNPs with more than 10% missing data across the 247 accessions and a minor allele frequency less than or equal to 1% were also filtered out. The imputation of missing data was performed using Beagle v5.2 (Browning et al., 2018) with default parameters and a genetic map as reported in Skovbjerg et al. (2023). A final set of 855,085 SNPs was used for GWAS analysis.

GWAS was conducted using three methods, namely MLM (Yu et al., 2006), FarmCPU (Liu et al., 2016), and mlmm.gwas (Segura et al., 2012). The Benjamini-Hochberg (BH) criterion was used for selecting a threshold p-value for each method and trait combination. When the threshold yielded no significant markers, a standard threshold of 10e^-5^ was used. Associated markers were visualised with a p-value distribution (expected vs. observed on a −log_10_ scale) with a Manhattan plot and a Q– Q plot. Each significant marker-trait association (MTA) was formatted and annotated with Planteome ontologies (Cooper et al., 2024), using the Plant Trait Ontology (Arnaud. et al., 2012), the Plant Ontology and the Plant Experimental Conditions Ontology. MTAs were integrated into the OrthoLegKB graph (Imbert et al., 2023), and connected to the marker-hosting genes. The query used to retrieve the GWAS results, presented in **Supplementary Table 1** is available in **Supplementary File 1**. Regions dense in MTAs were defined by clustering overlapping MTAs using a 675,000 bp interval on each side of the marker. A threshold of at least five MTAs was used to filter these regions.

### Gene functional annotation

Functional annotations including MapMan annotation (Thimm et al., 2004; Schwacke et al., 2019), Gene Ontology (GO) annotation (The Gene Ontology Consortium et al., 2023), and InterPro annotation (Jones et al., 2014; Blum et al., 2021) were performed for *P. sativum*, *V. faba,* and *L. culinaris* genes as described in Imbert et al., (2023). Annotations were queried in OrthoLegKB for each gene using the unique gene ID. An example of querying such information is available in **Supplementary File 1**.

### Differential gene expression analysis

Twenty-four *P. sativum* RNA-seq samples (NCBI BioProject PRJNA543764), corresponding to 2 varieties × 2 treatments × 3 sampling times × 2 biological replicates, from Bahrman et al. (2019), were used in this study. The two varieties from which aerial parts were sampled are Champagne which is a frost-tolerant accession and Térèse which is a frost-susceptible accession. Raw pseudo-counts for each RNA-seq sample were generated using Salmon (Patro et al., 2017), implemented in the nf-core/rnaseq pipeline v3.12.0 (Patel et al., 2023), converted to integers, and then subjected to differential expression analysis using the DESeq2 R package v1.42.1 (Love et al., 2014). Each sample was annotated using Planteome Plant Ontology (PO) and Plant Experimental Conditions ontology (PECO) and the pseudo-counts for each gene integrated into OrthoLegKB. Data were submitted to multifactorial analysis using DESeq2 in order to highlight (1) differentially expressed genes under low non negative temperature (LT) within each genetic background considered and (2) differentially expressed genes under LT or control conditions when comparing Champagne and Térèse accessions. Differentially expressed genes with an adjusted *p*-value < 0.05 and a | log2 fold change | > 1 were retained for further analysis.

The same procedure as described above was used to analyse *C. arietinum* RNA-seq samples from Akbari et al. (2023), using the frost-tolerant line Saral and frost-susceptible line ILC533 (available in NCBI BioProjects PRJNA905065 and PRJNA903665). The queries to retrieve transcriptomic data from OrthoLegKB are available in **Supplementary File 2**.

### Collection of QTL for frost tolerance, and for phenological, physiological and morphological traits

The scientific literature was searched for QTL related to frost tolerance, phenological such as flowering, physiological and morphological traits. Molecular markers were primarily aligned on the species respective reference genomes with minimap2 (Li, 2018) or with BLAT (Kent, 2002) and BLAST (Altschul et al., 1990) in case context sequences were PCR primers. For frost tolerance, this search allowed the retrieval of 14 QTL from Carrillo-Perdomo et al. (2022) from Hiverna x Silian (POP2) and Hiverna x Quasar (POP3) recombinant inbred line (RIL) populations, 22 QTL from Sallam et al. (2016a, 2022) in *V. faba* from the GWBP, 70 QTL from Beji et al. (2020) from a *P. sativum* diversity panel, 32 QTL from Boutet et al. (2023) from Champagne x Térèse RIL population, 16 QTL from Klein et al. (2014) from China x ‘Caméor’ *P. sativum* and 12 QTL from Avia et al. (2013) in *M. truncatula* from F83005-5 x DZA045-5 LR3 RIL population. In order to simultaneously query QTL for different variables of the same more general trait, all QTL were annotated with ontologies prior to their integration into OrthoLegKB. For instance, the TO (Cooper et al., 2024) general term TO:0000303 (low temperature stress response) for cold tolerance was connected to both frost damage (TO:0001152; frost damage stress response) and winter survival (TO:0001153; survival rate after freezing temperature stress) terms in this study. For the multi-trait analysis, QTL with the following annotations were considered: cold tolerance, flowering time, carbohydrate content, fat and essential oil content, plant axis morphology, phyllome morphology, whole plant morphology, and cardinal organ part morphology.

### Identification of orthologous genes and syntenic QTL

The Ortho_KB structural pipeline (Imbert et al., 2023), developed to compute homology between genes and synteny across chromosomal regions, was used to highlight orthologous genes and syntenic regions. Syntenic regions were here defined as regions showing conserved gene content and order. Orthologous genes and syntenic regions were then queried as shown in **Supplementary File 3**. Genes of the same OrthoFinder (Emms and Kelly, 2019) orthogroup located in syntenic blocks in different cool-season legumes revealed by MCScanX (Wang et al., 2012) were considered syntenic orthologous genes, referred to as syntenologs in this study. OrthoFinder v2.5.5 was run with its default parameters, using Diamond in ultra-sensitive mode for the alignment (Buchfink et al., 2021). The following parameters were used to run MCScanX: -k 50, -g −1, -s 10, -e 1e^-05^, -m 25, -w 5. To extract syntenic ortholog sets across more than two species, orthology and synteny relationships were first collected from OrthoLegKB for pairs of syntenologs, and then sets of syntenologs across species were detected by identifying cliques in a graph, i.e. fully connected groups of syntenologs, using the R library igraph v2.0.3 (Csardi and Nepusz, 2006). A threshold of three syntenic orthologous genes was applied to detect cliques.

## Results

### Genome-wide analysis of frost tolerance in *V. faba*

A *V. faba* diversity panel of 247 accessions was sown in autumn in three locations in France and two years, making a total of four environments: Bretenière and Theix in 2013-2014 and Bretenière and Orsonville in 2016-2017 (**Fig. 1**). Bretenière 2013-2014 registered 37 days with negative temperatures (below 0°C) versus 66 days in 2016-2017. Theix 2013-2014 and Orsonville 2016-2017 recorded respectively 85 and 48 days of negative temperatures over the course of the trials. The intensity of the frost was also different across environments with registered minimal temperatures of −4.7°C, −9.6°C, −11.4°C and −10.4°C in Bretenière 2013-2014 and 2016-2017, Theix 2013-2014 and Orsonville 2016-2017, respectively.

**Figure 1.**
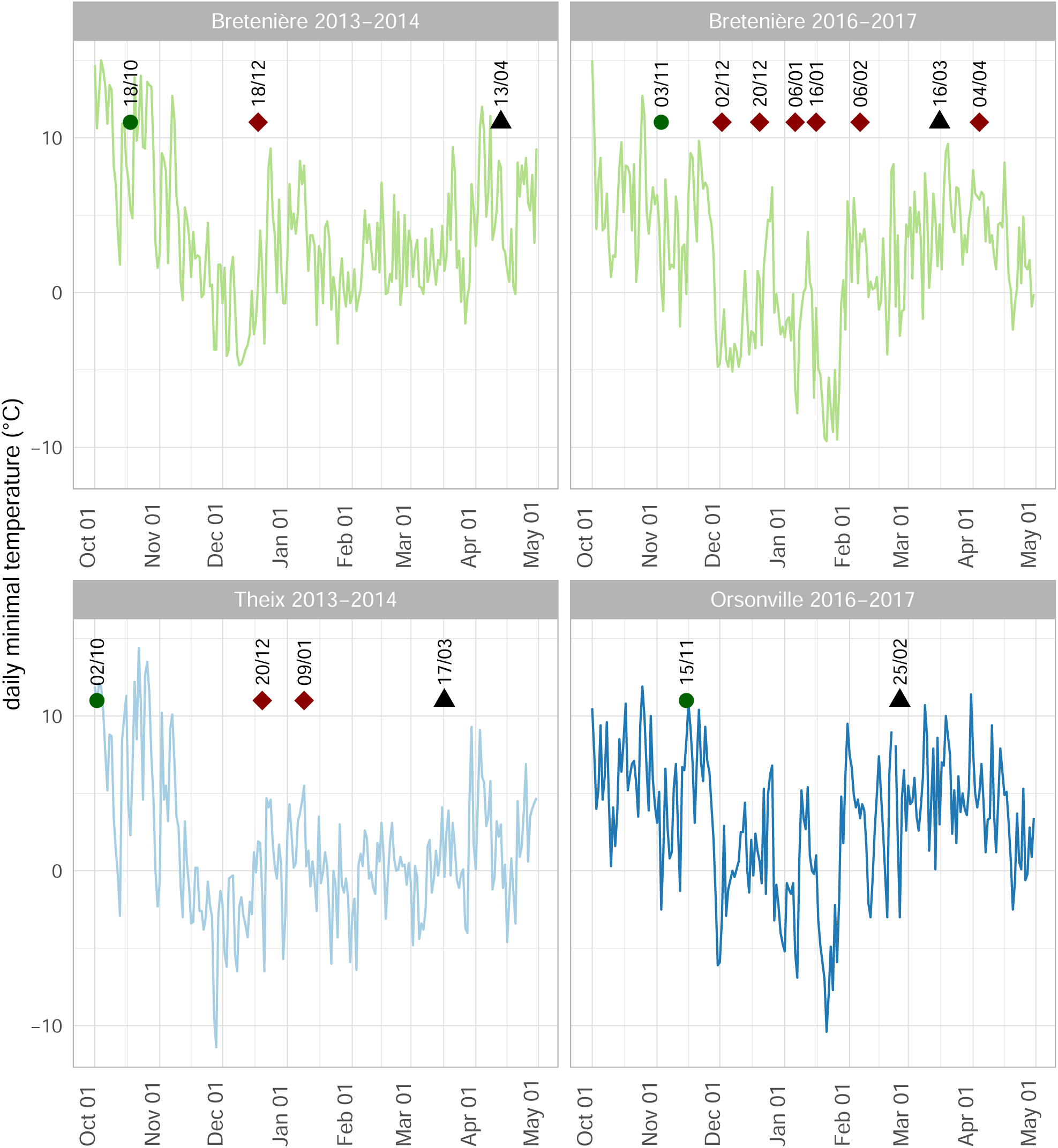
Minimum daily temperatures recorded under shelters at experimental sites. Sowing dates are indicated by green dots. Plants were phenotyped for frost damage after major frost events. Frost damage phenotyping dates are indicated by red diamonds above the curves. Plant survival was assessed at the end of the winter season. The phenotyping dates are indicated by dark triangles.

A GWAS analysis considering frost damage and winter survival scores from the four environments and using 855,085 exome capture-based markers mapped to the latest *V. faba* genome assembly (Jayakodi et al., 2023) revealed a total of 670 significant Marker-Trait Associations (MTAs; **Supplementary Table 1**). These 670 MTAs correspond to 405 unique markers, of which 303 were associated with frost damage and 143 with winter survival. Forty-one markers were associated with both traits, and 20 markers were detected by all methods, namely MLM, mlmm.gwas and FarmCPU (**Fig. 2a**). Of the 405 significant markers, 100 markers were identified for Bretenière in 2013-2014, 185 for Bretenière in 2016-2017, 112 for Theix in 2013-2014 and 28 for Orsonville in 2016-2017 (**Table 1**). MTAs were detected on all *V. faba* chromosomes, with 92 significant markers on Vf01S, 52 on Vf01L, 77 on Vf02, 44 on Vf03, 53 on Vf04, 62 on Vf05 and 25 on Vf06 (**Fig. 2b**). And, 18 markers were significant in at least two environments (**Fig. 2b**) including two markers significant in three environments, Vf01S:676945582 and Vf01S:676945977. These two markers were associated with frost damage and winter survival in Bretenière 2016-2017, in Theix 2013-2014 and in Orsonville 2016-2017. However, no significant association was consistently detected in all four environments.

**Figure 2.**
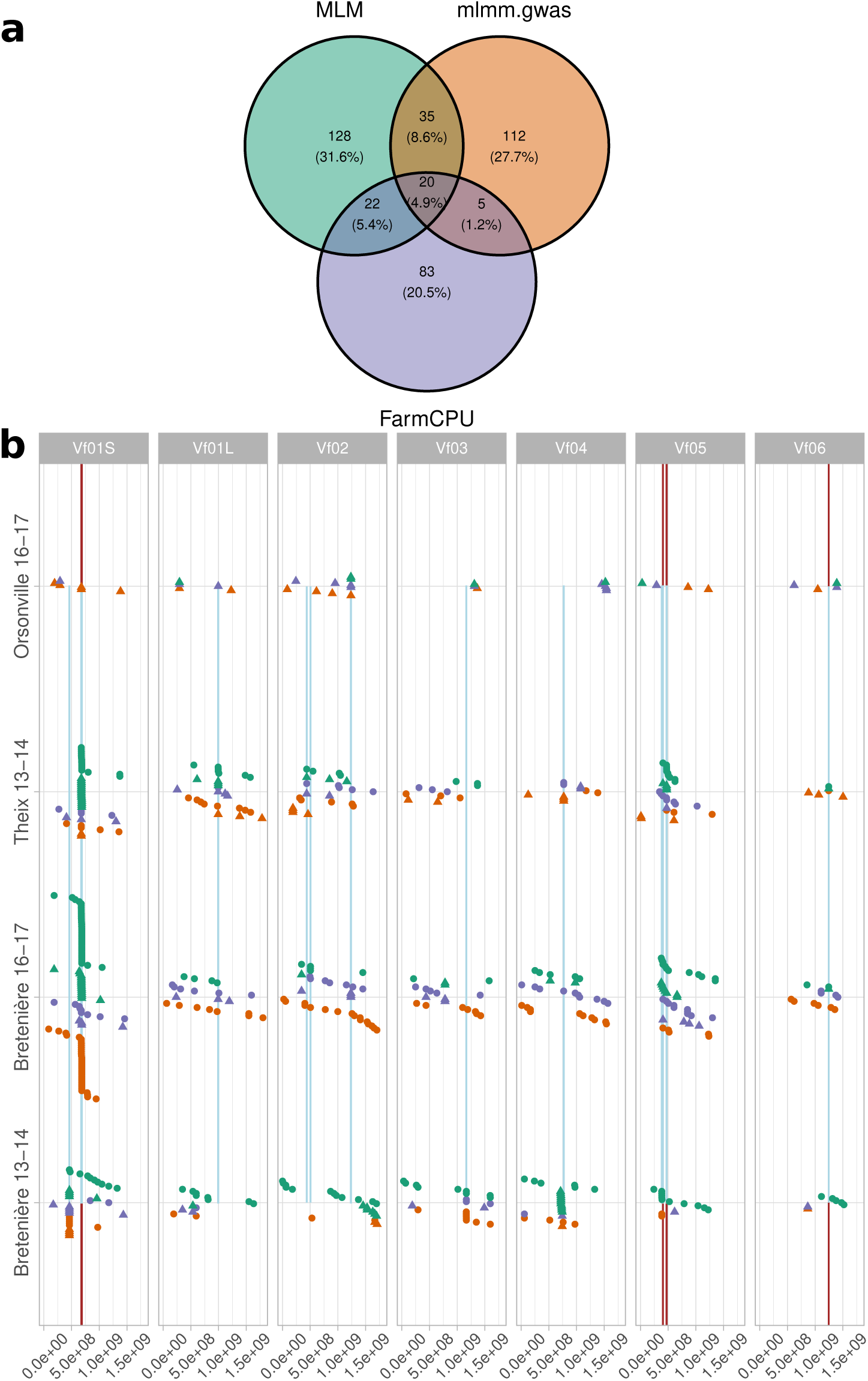
Overview of the significant marker-trait associations (MTAs) highlighted by genome-wide association study (GWAS) for frost damage and winter survival in *Vicia faba*. a) Venn diagram of the statistically significant SNP markers identified by each GWAS method, b) Genomic locations, by chromosome, of significant markers revealed using data from each environment. Markers associated with frost damage and survival rate traits are represented by dots and triangles. The colour of the shape indicates the method that allowed the detection of the association, with green, violet and orange referring to MLM, FarmCPU and mlmm.gwas, respectively. Vertical lines in brown and segments in light blue indicate significant markers identified in at least two environments and regions dense in MTAs, respectively.

**Table 1.**
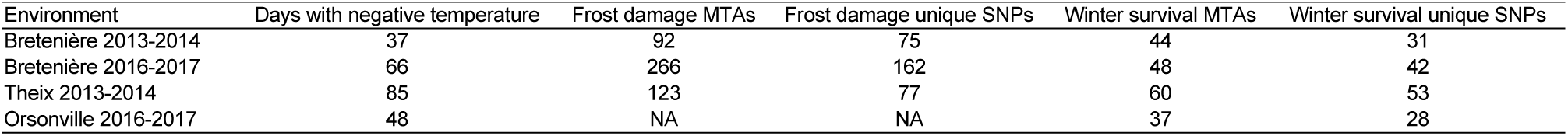
Number of marker-trait associations and unique markers identified across environments.

Nineteen genomic regions with a high density of significant markers were identified using a linkage disequilibrium-based approach. These regions were revealed by markers identified either by multiple methods, or in multiple environments, or both (**Supplementary Table 2**). The regions are found on all chromosomes of *V. faba*, as depicted in **Fig. 2b**. Fourteen of the 19 loci were associated with both frost damage and winter survival and detected in at least two environments, which are described hereafter. The 340-kb interval on Vf01S contained a putative organic cation/carnitine transporter-encoding gene, *Vfaba.Hedin2.R1.1g070920*, and another gene of unknown function. A 9-Mb genomic region of Vf01S was also detected. This region encompassed 179 MTAs and harboured four *CBF/DREB1* genes, *Vfaba.Hedin2.R1.1g099680*, *Vfaba.Hedin2.R1.1g099880*, *Vfaba.Hedin2.R1.1g100280*, *Vfaba.Hedin2.R1.1g101280*. The region also included *Vfaba.Hedin2.R1.1g099440* encoding an ATP-dependent Clp protease, *Vfaba.Hedin2.R1.1g099480* encoding an ABCB transporter, *Vfaba.Hedin2.R1.1g099840* encoding a glycosylphosphatidylinositol (GPI)-specific phospholipase, and *Vfaba.Hedin2.R1.1g100880* encoding a TBP-associated factor 8. On V01L, a 2-Mb region included seven MTAs and four genes, namely *Vfaba.Hedin2.R1.1g352040* encoding a phospholipase-D, *Vfaba.Hedin2.R1.1g352000* encoding a SR45a splicing factor, *Vfaba.Hedin2.R1.1g351640* encoding a sulfurtransferase and *Vfaba.Hedin2.R1.1g352200* encoding a RNA polymerase II large subunit. On Vf02, a 1-Mb locus of 8 MTAs contained *Vfaba.Hedin2.R1.2g194440* encoding a putative peptidyl-tRNA hydrolase and one gene encoding an uncharacterised protein. The genomic locus of 1-Mb on Vf04 encompassed six MTAs, and genes *Vfaba.Hedin2.R1.4g116520*, *Vfaba.Hedin2.R1.4g116800*, *Vfaba.Hedin2.R1.4g116840* and *Vfaba.Hedin2.R1.4g116920* encoding a phosphoinositide transfer protein, a member of the cytochrome P450 family, a protein belonging to the uncharacterised UPF0261 protein family and a kinetochore protein, respectively. On Vf05, a first 491-kb region of nine MTAs included *Vfaba.Hedin2.R1.5g063200* encoding an ADP binding protein and a gene encoding a protein of unknown function. A second region spanning 63 Mb and 13 MTAs included *Vfaba.Hedin2.R1.5g073520* encoding a peptidase and *Vfaba.Hedin2.R1.5g074200* encoding a lecithin:cholesterol acyltransferase. Finally, on Vf06, the identified region encompassed a single gene supported by seven MTAs, *Vfaba.Hedin2.R1.6g160560*, encoding a membrane attack complex/perforin (MACPF) domain-containing protein.

The candidate genes identified in this section, along with those presented in the subsequent sections, are listed in **Table 2**, along with supporting evidence.

**Table 2.**
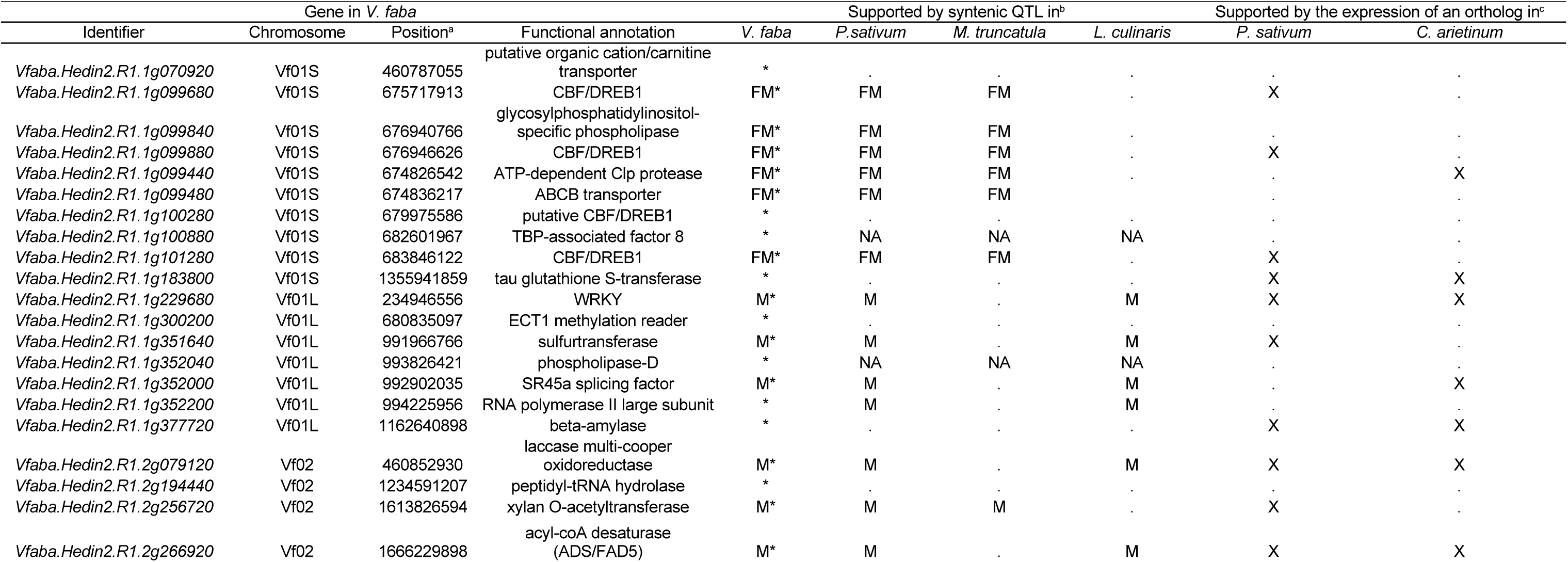
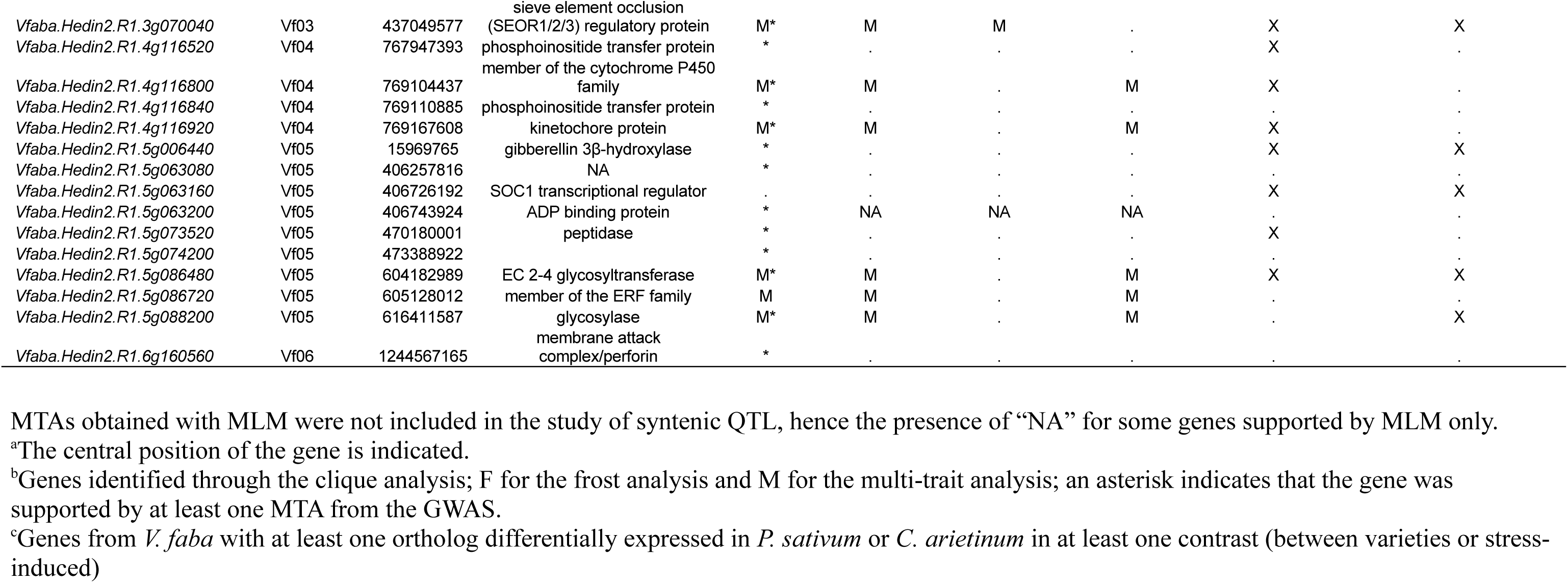
List of candidate genes for frost tolerance in *Vicia faba*. For each gene, its position on the corresponding *V. faba* chromosome, and functional annotation is indicated. Information supporting the gene as a candidate, either genetic or expression are also provided.

### Identification of syntenic frost tolerance QTL among cool-season legumes

We searched for synteny between frost tolerance QTL in *V. faba* and in *P. sativum* and *M. truncatula* because (1) these species are all native of temperate regions and therefore are likely to have evolved similar adaptations to frost, (2) their genomes are highly syntenic (**Supplementary Figs. 2 and 3** and **Supplementary Table 3**), and (3) genetic bases for frost tolerance have been reported for all three species. As the MLM model generates a larger number of significant markers than FarmCPU and mlmm.gwas, we decided to further consider only the MTAs supported by FarmCPU and mlmm.gwas in this section and in the following. This reduced the total number of *V. faba* MTAs from 670 to 353, as shown in **Supplementary Fig. 1a**. MTAs were annotated and the information were integrated into OrthoLegKB along with other public data and data originating from *M. truncatula* and *P. sativum* bringing the total number of QTL in the database to 520 (**Table 3; Supplementary Fig. 4a**).

**Table 3.**
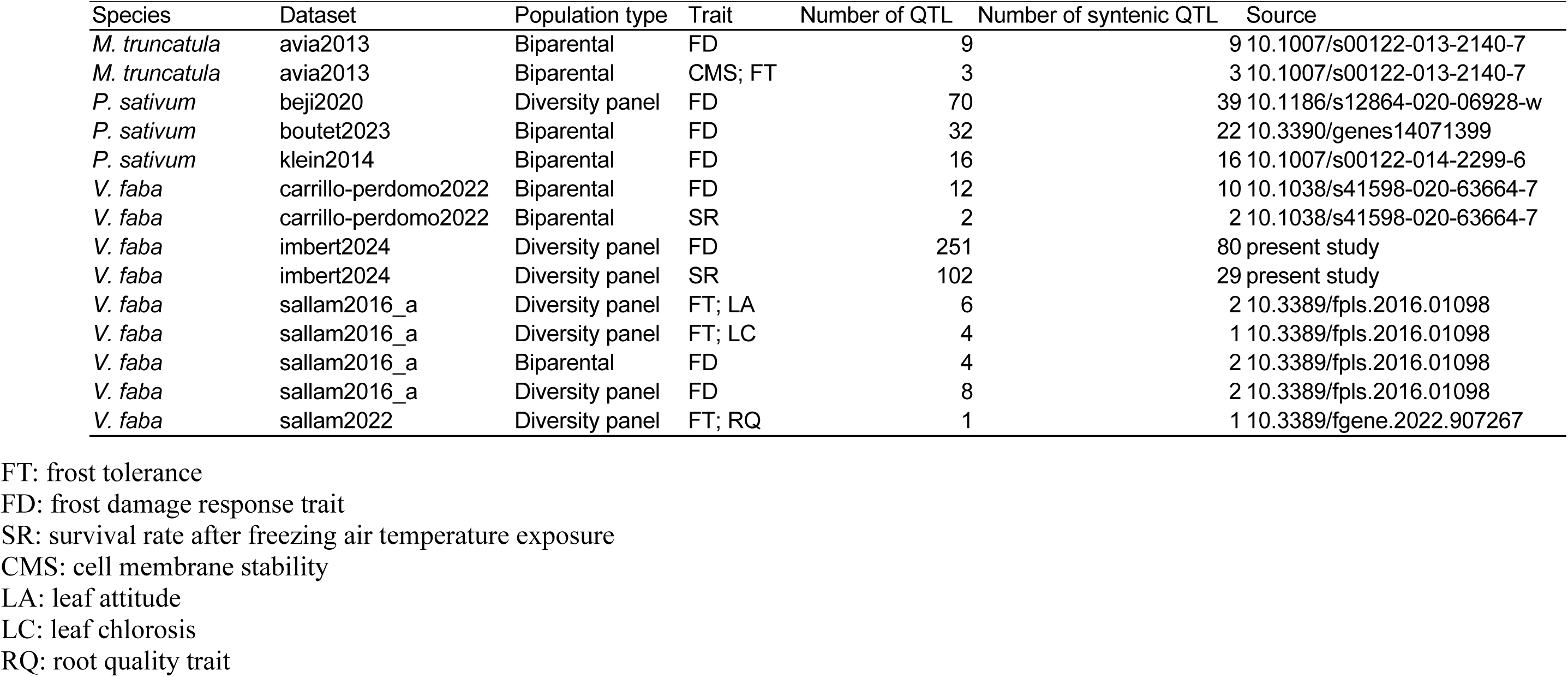
Overview of frost tolerance QTL integrated in OrthoLegKB along with the number of QTL found syntenic.

Of the 353 *V. faba* MTAs, representing a unique number of 277 significant markers, 109 significant markers, linked to 63 genes and located on Vf01S, Vf02, Vf03, Vf05 and Vf06, had syntenic connections with QTL for frost tolerance in other species. A set of 218 syntenic QTL for frost tolerance was retrieved from OrthoLegKB (**Supplementary Fig. 4a**). **Figure 3** shows the synteny between MTA loci, 12 QTL from Carrillo-Perdomo et al. (2022) and eight QTL from Sallam et al. (2016a, 2022) in *V. faba*, 39 QTL from Beji et al. (2020), 22 QTL from Boutet et al. (2023) and 16 QTL from Klein et al. (2014) in *P. sativum* and 12 QTL from Avia et al. (2013) in *M. truncatula* (**Supplementary Table 4**).

**Figure 3.**
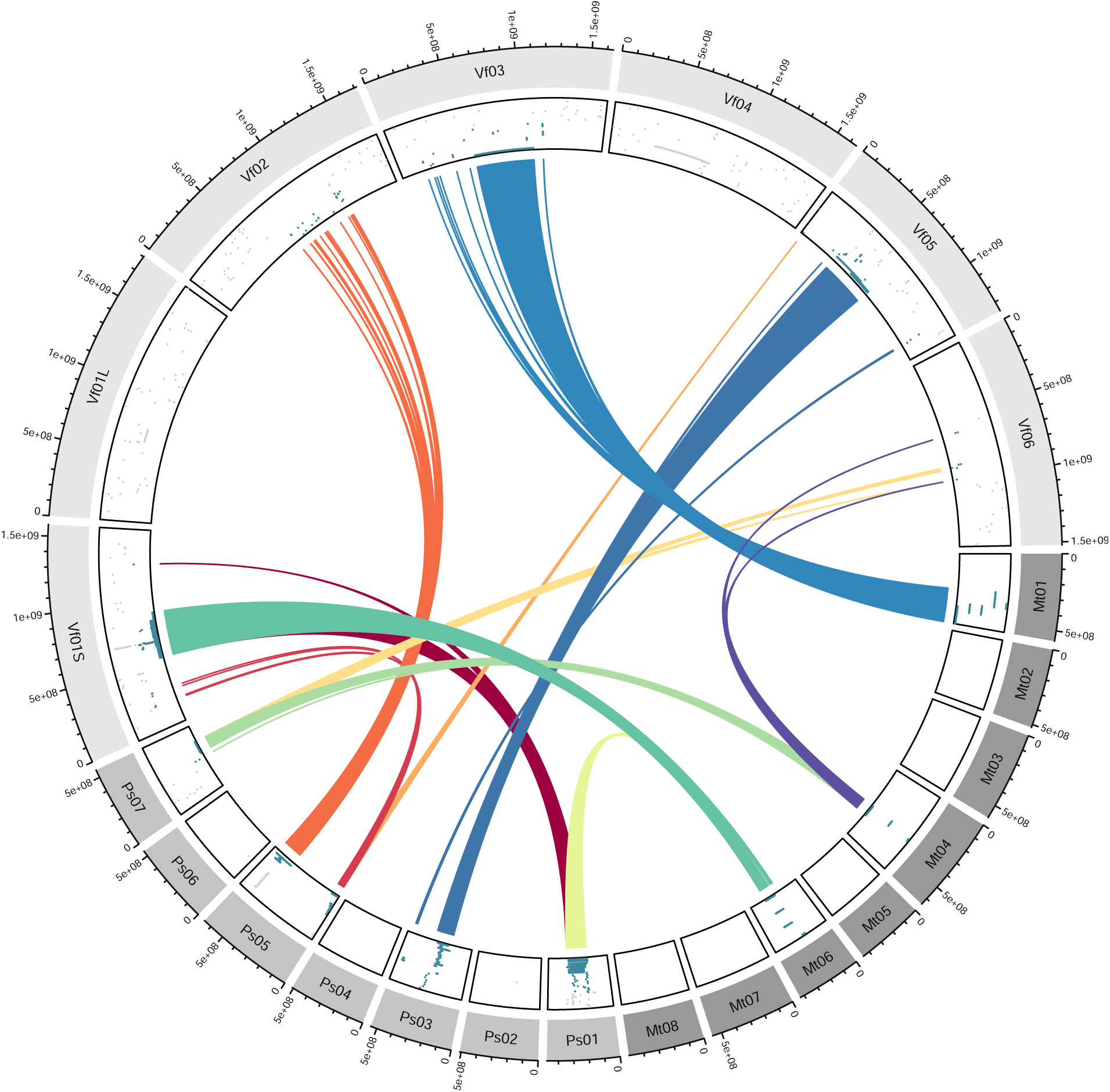
Syntenic frost tolerance QTL from *Vicia faba*, *Pisum sativum* and *Medicago truncatula*. The outer layer of the Circos corresponds to the chromosomes of all three species and the inner layer shows the positions of the QTL for frost tolerance from this study or from equivalent approaches carried out earlier and reported in the literature. Syntenic QTL are depicted in blue while non-syntenic QTL are in gray. Coloured ribbons connect syntenic QTL. QTL shown in this figure are either from QTL mapping in biparental populations or from GWAS and have all been integrated into OrthoLegKB. *M. truncatula* chromosome sizes have been increased by a factor of 10 for readability.

Of all syntenic regions harbouring QTL for frost tolerance, those on *V. faba* Vf01S, *P. sativum* Ps01, *M. truncatula* Mt06, on the one hand, and Vf06, Ps07 and Mt04 on the other hand were particularly conserved across the three species (**Fig. 3**). Other *V. faba* MTA loci had syntenic counterparts in only one species. For example, frost tolerance QTL on Vf02, Vf03, and Vf05, were found to be in syntenic regions to those hosting frost tolerance QTL on Ps05, Mt01, and Ps03, respectively (**Fig. 3**).

Some of the QTL regions were supported by a limited number of QTL. For example, the conserved syntenic region between the three species on Vf06, Ps07, and Mt04 was supported by only one large QTL on Ps07 from Klein et al. (2014), while six MTAs were identified in five genes on Vf06 and three QTL were reported on Mt04 (Avia et al., 2013). Conversely, relatively more individual QTL were identified in the conserved region on Vf01S, Ps01 and Mt06.

### Colocalisation of syntenic frost tolerance QTL on *Vicia faba* Vf01S, *Pisum sativum* Ps01 and *Medicago truncatula* Mt06 with *CBF/DREB1* gene clusters

Genomic regions harbouring QTL for frost tolerance on *V. faba* Vf01S, *P. sativum* Ps01 and *M. truncatula* Mt06 (**Fig. 3**) contain large clusters of *CBF/DREB1* genes. The regions were annotated with 14, 10, and 12 genes in the *V. faba* Hedin/2 v1.0 (Jayakodi et al., 2023), *P. sativum* ‘Caméor’ v2 (in prep), and *M. truncatula* A17 v5.0 (Pecrix et al., 2018) genome assemblies, respectively (**Fig. 4a**). Except for *V. faba Vfaba.Hedin2.R1.1g100280* and *P. sativum Psat.cameor.v2.1g160450* genes that had exceptional large sizes, the *CBF/DREB1* genes from the syntenic regions fell into the same orthogroups but no simple orthologous relationships could be revealed at the interspecific level (**Fig. 4b**).

**Figure 4.**
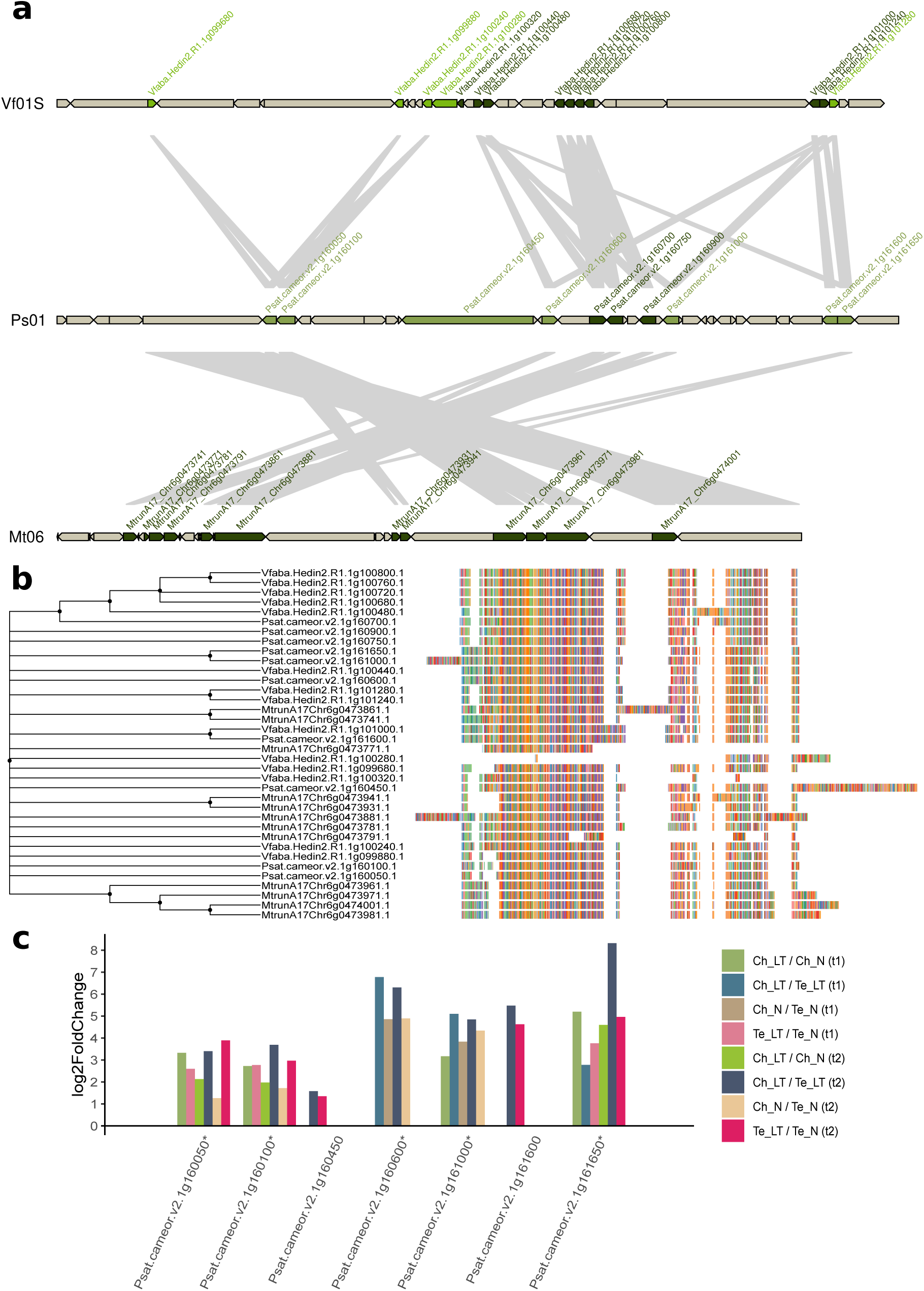
Composition of and synteny between *CBF/DREB1*-rich genomic regions on *Vicia faba* Vf01S, *Pisum sativum* Ps01 and *Medicago truncatula* Mt06. a) Microsynteny in the *CBF/DREB1* loci colocating with cold tolerance syntenic QTL in *V. faba*, *P. sativum* and *M. truncatula*. Genes are represented by arrows indicating the orientation of the open reading frames, and *CBF/DREB1* genes are highlighted in different shades of green. Ribbons connect *CBF/DREB1* gene pairs belonging to the same orthogroup (Emms and Kelly, 2019). *V. faba CBF/DREB1* genes harbouring SNP markers significantly associated with frost tolerance-related traits in our GWAS are in light green. *P. sativum CBF/DREB1* genes identified as differentially expressed in response to low temperature or constitutively differentially accumulated in the tolerant cultivar are in khaki. Other *CBF/DREB1* genes are shown in dark green. As the three species have high heterogeneity in genome size and variable intergenic sizes, intergenic regions are not depicted. However, the gene sizes remain proportional. The genomic intervals illustrated in this figure are: *V. faba* v1.0 (Jayakodi et al., 2023) Vf01S:675717602-683846462 bp, *P. sativum* v2 (‘Caméor’ v2 in prep.; for v1, see Kreplak et al., 2019) Ps01:196133118-199952977 bp, and *M. truncatula* v5.0 Mt06:27136282-27451503 bp (Pecrix et al., 2018). Notably, although not shown here, *Psat.cameor.v2.1g161600* actually has an ortholog in *M. truncatula*, *MtrunA17_Chr6g0484671*, which is located further away from the main cluster. b) Phylogenetic analysis of *CBF/DREB1* genes contained in the regions of interest. The phylogenetic consensus tree was generated using protein sequences, the maximum likelihood method and Whelan and Goldman model over 1,000 bootstrap replicates in MEGAX v10.2.6. The alignments obtained with MUSCLE (Edgar, 2004) and used to construct the phylogenetic tree are presented on the right side of the tree. c) Fold-change of putative *CBF/DREB1 P. sativum* genes of the locus under consideration, found differentially expressed in at least one condition. Genes with orthologs in *V. faba* bearing markers significantly associated with frost tolerance are indicated with asterisks. Ch: Champagne; Te: Térèse; N: control; LT: low temperatures; T1: sampling time point 1; T2: sampling time point 2.

To support the putative role of these genes in response to low temperature (LT), we attempted to use available transcriptomic resources and in particular the study by Bahrman et al. (2019), which was published prior to the release of the first genome assembly of *P. sativum*. Briefly, an RNA-seq dataset derived from the aerial parts of young *P. sativum* plants was used to reveal molecular determinants of response to LT treatment in two accessions contrasted for frost tolerance. Champagne is known to be frost tolerant while Térèse is frost susceptible (Bahrman et al., 2019). We performed a differential expression analysis to reveal up- and downregulated genes either in Champagne comparatively to Térèse or under LT comparatively to control conditions (**Supplementary Table 5; Supplementary Material 1**). Evidence was found that genes belonging to the same orthogroup may have similar functions. Indeed, *Vfaba.Hedin2.R1.1g099680, Vfaba.Hedin2.R1.1g099880*, and *Vfaba.Hedin2.R1.1g100240* were in the same orthogroup as *Psat.cameor.v2.1g160050* and *Psat.cameor.v2.1g160100*. All three *V. faba* genes hosted markers significantly associated with frost tolerance and both *P. sativum* orthologs were differentially expressed under LT (**Fig. 4c**). Moreover, genes *Psat.cameor.v2.1g160600, Psat.cameor.v2.1g161000* were upregulated only in Champagne under LT (**Supplementary Table 5**). In contrary, genes *Vfaba.Hedin2.R1.1g100480, Vfaba.Hedin2.R1.1g100680, Vfaba.Hedin2.R1.1g100720, Vfaba.Hedin2.R1.1g100760, Vfaba.Hedin2.R1.1g100800, Psat.cameor.v2.1g160700, Psat.cameor.v2.1g160750,* and *Psat.cameor.v2.1g160900,* all part of the same orthogroup, did neither host MTAs nor were found responsive to LT (**Fig. 4**).

### Synteny between frost tolerance QTL on *Vicia faba* Vf05 and frost tolerance QTL on *Pisum sativum* Ps03 spanning Mendel’s *Le* locus

Given that the *V. faba* chromosome Vf05 is largely syntenic with the *P. sativum* chromosome Ps03, we were intrigued by the identification of a significant MTA from the present study at the top of Vf05, which belonged to a region syntenic with the top of Ps05. The syntenic region connecting the tops of Ps05 and Vf05 over 21.5 and 47.8 Mb, contained multiple QTL for frost tolerance derived from either field or controlled environment experiments (Klein et al., 2014; Beji et al., 2020; Boutet et al., 2023) and 426 and 273 genes, respectively (**Supplementary Fig. 5**). The MTA locus in question, on Vf05, associated with winter survival, was revealed by mlmm.gwas in Theix in 2013-2014. The marker was located in *Vfaba.Hedin2.R1.5g006440*, the ortholog of *Psat.cameor.v2.5g10750* or *PsGA3ox1*, also known as Mendel’s stem length gene (*Le*) (**Supplementary File 4**; Lester et al., 1997), which encodes a gibberellin 3β-hydroxylase and controls internode elongation. *PsGA3ox1* was also revealed by a MTA for frost tolerance in the vicinity of the gene (Beji et al., 2020). The expression level of *PsGA3ox1* was significantly reduced when the *P. sativum* varieties Champagne and Térèse were subjected to LT (**Supplementary Table 5**). Indeed, a significantly lower expression was observed in the frost tolerant accession Champagne compared to Térèse. Transcriptomic evidence in *C. arietinum* from Akbari et al. (2023) showed that this gene is also upregulated in response to LT in cold-tolerant Saral and cold-susceptible ILC533 lines (**Supplementary Table 6**).

### Conserved syntenic orthologs under frost tolerance QTL: genetic, genomic and transcriptomic evidence

Syntenic orthologous genes, hereafter called syntenologs, underlying the frost tolerance QTL in *V. faba*, *P. sativum*, and *M. truncatula* were searched for, using OrthoLegKB. A total of 26 orthogroups with orthologs from all three species being all connected to frost tolerance QTL was identified (**Supplementary Table 7**). Notably, seven orthogroups contained genes encoding transcription factors; three encoding CBF/DREB1 transcription factors, while the other four orthogroups included genes encoding ARR-B-type, bHLH class-Ib, CCCH zinc finger and MADS/AGL transcription factors respectively. *CBF/DREB1* genes all belonged to the syntenic region described in **Fig. 4**. The *P. sativum* gene encoding a MADS/AGL transcription factor, *Psat.cameor.v2.1g157700*, was upregulated in Champagne under both controlled and LT conditions. Similarly, *Psat.cameor.v2.1g156050*, encoding a bHLH class-Ib was more expressed in Champagne compared to Térèse under LT. We also found genes encoding proteins other than transcription factors, which were differentially expressed in *P. sativum*. For instance, genes *Psat.cameor.v2.1g157600* encoding a gibberellin 20-oxidase and *Psat.cameor.v2.1g157800* encoding a fructose kinase showed a significantly lower expression under control conditions in Champagne comparatively to Térèse. As an other example, the gene *Psat.cameor.v2.1g163150* encoding for a BAG nucleotide exchange factor was downregulated in Champagne under LT. Some other orthogroups included one-to-one syntenologs that were not differentially expressed in *P. sativum* (**Supplementary Table 5 and Supplementary Table 7**).

### Identification of syntenic regions controlling multiple traits, including frost tolerance

Synteny between frost tolerance controlling QTL (**Supplementary Table 4)** and QTL controlling plant morphology, phenology and physiology (**Supplementary Fig. 4b**) was investigated. Details on the queried QTL integrated into OrthoLegKB are given in **Supplementary Table 8**.

First, considering only *V. faba* data, we searched for QTL co-localising with MTA loci. Although no QTL for other traits were identified to colocalise with the main loci detected on Vf01S and Vf05 (**Fig. 5**), *V. faba* chromosome Vf01L was found to carry two QTL for frost damage and a QTL for linoleic acid content in cold-acclimated plants from Sallam et al. (2016a) in a region of 92.6 Mb. This interval also contained a MTA from the present study, located in the gene *Vfaba.Hedin2.R1.1g300200*, which encodes an ECT1 methylation reader.

**Figure 5.**
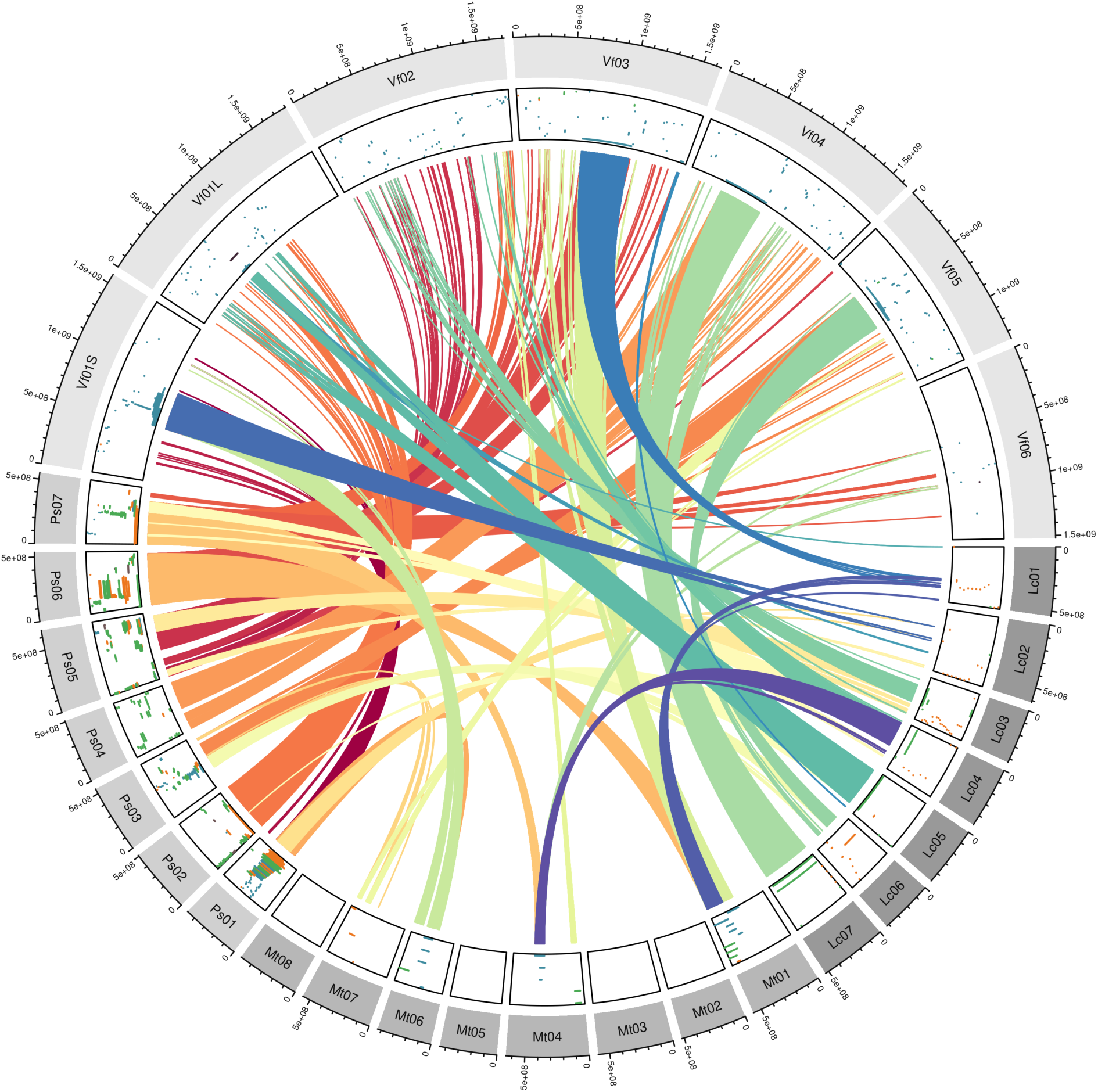
Overview of frost tolerance, morphological, phenological and physiological QTL syntenic with frost tolerance QTL. The considered species are *Vicia faba*, *Pisum sativum*, *Medicago truncatula* and *Lens culinaris*. The outer layer of the circos corresponds to the chromosomes of all four species and the inner layer shows the positions of the QTL from this study or from equivalent approaches carried out previously and reported in the literature. QTL for frost tolerance, morphological, phenological and physiological traits are in blue, green, orange and dark brown, respectively. Coloured ribbons connect syntenic QTL. QTL shown in this figure are either from QTL mapping approaches in biparental populations or from GWAS and have all been integrated into OrthoLegKB. *M. truncatula* chromosome sizes have been increased by a factor of 10 for readability.

Then, we searched in other cool-season legume species for genomic intervals identified as controlling frost tolerance that also included QTL for other traits. Multiple colocalisations were highlighted as illustrated in **Fig. 5**. For example, in *P. sativum,* a total of 27 QTL encompassing the *CBF/DREB1* gene-rich region on Ps01 were identified. These QTL control days to flowering, shoot branching, shoot dry weight, stipule length and leaf area index. In addition, 24 colocalising QTL were identified in the *Le* region on *P. sativum* Ps05 including QTL for flowering, leaf chlorosis under *Aphanomyces euteiches* infestation, plant height and axis morphology (**Supplementary Table 9**).

When extending the analysis across species, we found that the syntenic frost tolerance loci on *P. sativum* Ps03 and *V. faba* Vf05 were also linked to QTL for morphological and phenological traits. Indeed, the genomic interval between 279 and 295 Mb on Ps03 included several *P. sativum* frost tolerance QTL from Beji et al. (2020), one QTL for branch number from Klein et al. (2014), as well as three flowering QTL from Williams et al. (2022) and Gali et al. (2018). Interestingly, the syntenic region on *L. culinaris* Lc06 was reported to host a large QTL for days to flowering from Haile et al. (2021) with *LcFTa1* (*Lcu.2RBY.6g043850*) as a potential candidate gene (Yuan et al. 2021). *TERMINAL FLOWER* or *FT* genes (*FTb, FTa2, FTc*) have not been detected by our GWAS. However, *Vfaba.Hedin2.R1.5g088200* and its syntenologs (**Supplementary Table 10**), encoding putative cell wall-localised GPI-anchored β-1,3 glucanases, are worth considering as candidate genes to explain at least frost tolerance. In fact, *Vfaba.Hedin2.R1.5g088200* hosted a marker significantly associated with frost tolerance in this study. We also identified two QTL for flowering from Klein et al. (2014) on *P. sativum* Ps07, syntenic with two QTL for frost tolerance from Avia et al. (2013) on *M. truncatula* Mt04, more specifically associated with the shoot dry weight and leaf area index under freezing conditions. At the syntenic locus on *V. faba* Vf02, a MTA for frost tolerance from this study was identified in the gene *Vfaba.Hedin2.R1.2g256720*, which encodes a xylan O-acetyltransferase.

We identified groups of syntenologs involving at least three species, including *V. faba.* Among these syntenologs, we retained only those that were associated with QTL, at least one of which was for frost tolerance. We identified 1,193 orthogroups (**Supplementary Table 10**). Among the *V. faba* genes, 73 were revealed by our GWAS, some of which have *P. sativum* and *C. arietinum* orthologs differentially expressed in response to LT in Bahrman et al. (2019) and Akbari et al. (2023), in the same direction (**Supplementary Table 7 and 11**). We illustrate these findings with five candidate genes. First, on Vf02, *Vfaba.Hedin2.R1.2g079120*, encoding a putative laccase multi-copper oxidoreductase, belong to a region syntenic to Ps05, which contains nine relatively large QTL for plant height from Pecrix et al. (2018) and Wu et al. (2021). The syntenic counterpart on Lc06 underlies a QTL for root sectional area QTL (Ma et al., 2020). Under LT, orthologs of this gene were downregulated in tolerant and susceptible lines in both species. Second, a WRKY transcription factor encoding gene, *Vfaba.Hedin2.R1.1g229680*, located on *V. faba* Vf01L, was also detected by our GWAS. In particular, its syntenologs were located in the confidence interval of the QTL for leaf chlorosis under *A. euteiches* attack (Leprévost et al., 2023) on Ps02 and for flowering (Yuan et al., 2021) on Lc05. The orthologs of this gene in *P. sativum* and *C. arietinum* were downregulated in susceptible lines under LT. Third, we detected genes encoding sieve element occlusion protein (SEO), which showed a higher expression in the tolerant varieties under LT compared to susceptible lines, and for which the *V. faba* ortholog *Vfaba.Hedin2.R1.3g070040* carried three MTAs from our study. The syntenologs on Mt01 and Ps06 were linked to a frost tolerance QTL from Avia et al. (2013) and an *A. euteiches* response leaf chlorosis QTL from Leprévost et al. (2023), respectively. Fourth, the gene *Vfaba.Hedin2.R1.2g266920* on Vf02, which encodes an acyl-coA desaturase (ADS/FAD5), showed syntenologs on Ps07 and Lc04 encompassed by QTL for flowering duration and time from Klein et al. (2014) and Neupane et al. (2022), respectively. Exposure to LT induced a lower expression level in both frost-tolerant lines in *P. sativum* and *C. arietinum*. Fifth, the gene *Vfaba.Hedin2.R1.5g086480* on Vf05, encoding an EC 2-4 glycosyltransferase carried an MTA from this study and was located in the interval of the frost tolerance QTL of Carrillo-Perdomo et al. (2022) and Sallam et al. (2016a). Its syntenologs on Ps03 and Lc06 fall in the confidence intervals of flowering time QTL from Gali et al. (2018) and Yuan et al. (2021), respectively. The orthologs of this gene were downregulated under LT in both species.

Interestingly, on Vf05, the gene *Vfaba.Hedin2.R1.5g086720*, encoding a member of the ERF family, was not found to be associated with frost tolerance in the present study, but was located under three frost damage QTL from Carrillo-Perdomo et al. (2022) and from Sallam et al. (2016a). It was also considered as a candidate gene for a plant height QTL from Skovbjerg et al. (2023). The region was also associated with flowering control, as the syntenologs on Ps03 and Lc06 underlie two QTL for this trait from Gali et al. (2018) and from Yuan et al. (2021), respectively (**Supplementary Table 10**).

## Discussion

### Genetic architecture of frost tolerance in *Vicia faba*

To identify the key genes controlling frost tolerance in *V. faba* and to promote the breeding of winter-hardy cultivars, we performed a GWAS on a diversity panel, including several frost-tolerant accessions originating from different countries, such as Hiverna from Germany (Link et al., 2010), Côte d’Or from France (Duc, 1997) and Quasar from the United Kingdom (Ondřej and Huňady, 2007). Accessions known to be frost-susceptible, such as Sillian from the Northern State of Sudan (Carrillo-Perdomo et al., 2022) were also included. We identified six robust genomic regions dense in MTAs for frost tolerance, located on chromosomes Vf01S, Vf02, Vf03 and Vf04. This is consistent with results obtained from biparental populations derived from crosses between Hiverna and Quasar or Hiverna and Sillian (Carrillo-Perdomo et al., 2022). These populations were evaluated for frost damage and survival rate in Bretenière in 2016-2017, a common environment with the diversity panel from this study. Carrillo-Perdomo et al. (2022) reported 14 frost tolerance QTL located on four linkage groups corresponding to *V. faba* chromosomes Vf01S, Vf03, Vf04 and Vf05. Out of these, ten QTL were located on chromosome Vf01S, in a *CBF/DREB1* gene-rich genomic region, which was also detected in the present study (**Fig. 2**). MTAs reported here also colocalised with the three QTL of Carrillo-Perdomo et al. (2022) on chromosomes Vf03, Vf04 and Vf05, confirming the robustness of the outcomes and supporting the candidate genes identified. Furthermore, the QTL on Vf05 from Carrillo-Perdomo et al. (2022) colocalized with a QTL for frost damage detected by Sallam et al. (2016a). This suggests that the interval located between 346,752,878 and 720,252,430 bp on Vf05 contains one or more robust genes involved in the control of frost tolerance.

With the exception of Vf01S and Vf05, few common regions were detected in multiple environments, suggesting that the effect of loci controlling frost tolerance is largely dependent on the specific winter conditions encountered (Arbaoui et al., 2008; Carrillo-Perdomo et al., 2022). Indeed, under field conditions, frost events may occur at different stages of the plant growing cycle, and with or without snow cover. In addition, plants may have reached different levels of hardening or dehardening by the time frost occurs. The plant response can also be influenced by various other stresses, including diseases, which can affect the overall performance during regrowth after the winter season (Boutet et al., 2023).

In addition to the regions shared with those of Carrillo-Perdomo et al. (2022) and/or Sallam et al. (2016a), we have identified new loci controlling frost tolerance on chromosomes Vf02 and Vf03, likely thanks to the wider genetic diversity considered and greater number of environments under which frost tolerance phenotyping was conducted. Further analysis of these loci will provide valuable insights to unravel their role in *V. faba* frost tolerance.

### Conserved genetic basis of frost tolerance across cool-season legumes

Comparative genomics enables the identification of conserved regions across species sharing similar blocks of genes inherited from a common ancestor, referred to as syntenic regions (Salse, 2022). Syntenic QTL found in different species, and especially for the same trait, may therefore indicate a shared genetic and molecular determinism. Thanks to the graph-based database OrthoLegKB (Imbert et al., 2023), we identified, in the present study, two sets of frost tolerance QTL being syntenic in the three legume species *V. faba*, *P. sativum* and *M. truncatula*.

Firstly, several MTAs were identified on *V. faba* Vf01S in a region containing a cluster of *CBF/DREB1* genes. Similar results were previously reported in *M. truncatula* (Tayeh et al., 2013b), and *P. sativum* (Tayeh et al., 2013b; Klein et al., 2014) and interestingly, QTL from *CBF/DREB1*-rich regions from the three species were found to be syntenic (**Fig. 4**). These findings corroborate with results from Carrillo-Perdomo et al. (2022) suggesting an important role for *CBF/DREB1* genes in controlling frost tolerance in cool-season legumes, similarly to what has been reported in non-legume species. Indeed, colocalisations between frost tolerance QTL and loci containing multiple copies of *CBF/DREB1* genes have been reported in multiple species, including *Triticum aestivum* (Sieber et al., 2016) and *A. thaliana* (Alonso-Blanco et al., 2005). Functional analyses have also contributed to demonstrate the role of *CBF/DREB1* genes in controlling the expression of a large number of genes responsible for frost tolerance (Zhao et al., 2016). Of the 14 putative *CBF/DREB1* genes annotated on Vf01S in *V. faba*, four genes hosted markers significantly associated with frost tolerance according to the three GWAS methods employed in our study (**Fig. 4**) and a fifth gene hosted markers which were only detected by the MLM method. Considering that the syntenic *CBF/DREB1* loci in *P. sativum* ‘Caméor’ and *M. truncatula* A17 contain 10 and 12 genes, respectively, the validation of the contribution of each *CBF/DREB1* gene to frost tolerance and a characterisation of the polymorphism, including copy number variation, at the inter- and intra-specific levels would deserve a detailed study in the future. In the absence of a publicly available transcriptomic dataset for frost tolerance in *V. faba*, we took advantage of the phylogenetic proximity between *V. faba* and *P. sativum* to assess the expression of *V. faba* orthologs. In agreement with the original paper from Bahrman et al. (2019), the reanalysis of the transcriptome samples using *P. sativum* cv. ‘Caméor’ genome assembly showed that, in *P. sativum*, some *CBF/DREB1* genes are differentially expressed between frost-tolerant and frost-susceptible genotypes constitutively and/or in response to LT. A similar study in *V. faba* would determine the extent of functional conservation of the *CBF/DREB1* genes.

Secondly, our results pinpointed the conservation of a major determinant of plant architecture across cool-season legumes. The *Le* locus in *P. sativum* is known to control internode length (Lester et al., 1997) and has been suggested to simultaneously control plant growth and frost tolerance (Burstin et al., 2007; Klein et al., 2014; Beji et al., 2020; Boutet et al., 2023). Here, we report synteny between *V. faba* and *P. sativum* at the *Le* locus (**Supplementary Fig. 5**). The *PsGA3ox1* gene (= *Le*), which encodes a Gibberellin (GA) 3β-hydroxylase that catalyses the conversion of GA_20_ to biologically active GA_1_ (Lester et al., 1997), has been reported to be responsible for the control of plant growth and development in *P. sativum* (Burstin et al., 2007). The observed reduction in *PsGA3ox1* expression level under LT, especially in the frost-tolerant genotype Champagne, is consistent with previous hypotheses of a morphological adaptation under LT conditions. While Champagne is significantly taller than Térèse under long days, this genotype exhibits a rosette-type dwarfism under short days in winter (Boutet et al., 2023). A similar rosette-type phenotype also exists in *V. faba* (Link et al., 2010). The link between cold acclimation and the gibberellin pathway has been described in *A. thaliana* (Achard et al., 2008). The short phenotype is suggested to create a microclimate providing a favourable effect on its response to frost and to biotic stresses (Körner, 2016; Boutet et al., 2023). It is therefore high likely that the ortholog of *PsGA3ox1* in *V. faba* does contribute to frost tolerance.

Finally, syntenic counterparts could not be detected for some frost tolerance QTL. This is the case for *V. faba* frost tolerance QTL identified on chromosomes Vf03 and Vf04 (Carrillo-Perdomo et al., 2022 and this study) or conversely for frost damage QTL located on chromosome Ps06 and Ps07 in *P. sativum* (Beji et al., 2020), that showed no syntenic QTL on the homologous chromosome Vf03 and Vf06 in *V. faba*, respectively. This can be attributed either to (1) a species-specific adaptation to frost, (2) some genomic rearrangements causing potential difficulties in detecting putative functional orthologs, (3) an absence of polymorphism in syntenic regions in the considered populations (Korte and Farlow, 2013), or (4) some differences in the experimental designs or environmental conditions of the different studies conducted. Future GWAS and QTL mapping studies, once conducted and integrated into OrthoLegKB, may provide a more complete insight into the genetic architecture of frost tolerance and the extent of QTL conservation across species.

### Key candidate genes controlling frost tolerance identified using a multi-species integrated approach

The combination of QTL, genomic and transcriptomic data herein allowed the identification of several interesting candidate genes for frost tolerance in cool-season legumes (**Table 2**), including genes involved in reactive oxygen species (ROS) scavenging as well as on sugar- or cell wall-related responses to cold.

ROS are known to accumulate in stressed plants (Atkinson and Urwin, 2012), including under LT (Guo et al., 2022). We identified *Vfaba.Hedin2.R1.6g160560*, which encodes a MACPF domain-containing protein, with a role in cell death and immunity in *A. thaliana*, and possibly in the positive regulation of ROS (Zhang et al., 2022). Homologous genes in *Gossypium hirsutum* were shown to be co-expressed with gene encoding WRKY and AP2/ERF-domain-containing transcription factors, under cold stress, and silencing of a MACPF gene improved cold tolerance in the same species (Chen et al., 2021). Glutathione S-transferases (GSTs) are isozymes that are essentially involved in the negative regulation of ROS (Gullner et al., 2018) while exhibiting a diversity of specific functions (Sappl et al., 2009). *Vfaba.Hedin2.R1.1g183800* encodes a putative member of the primary class tau of the superfamily of GSTs and has three orthologs in *P. sativum*. The three *P. sativum* orthologs were upregulated in the frost-tolerant genotype Champagne under LT (**Supplementary Material 1**), as highlighted by the reanalysis of the gene expression dataset of Bahrman et al. (2019). A GST class phi has been shown to interact with INDUCER OF CBF EXPRESSION 1 (ICE1) which is thought to induce the expression of *CBF/DREB1* genes *(Gilmour et al., 1998; Thomashow and Torii, 2020; Shen et al., 2024)*.

The accumulation of sugars participates in the response to LT as osmolytes, but especially through the stabilisation of plasma and organelle membranes. Notably, raffinose and maltose are involved in this function, in ROS scavenging and in the protection of photosystems (Tarkowski and Van den Ende, 2015). Indeed, raffinose is frequently cited in the literature as a soluble sugar that accumulates in legumes in response to LT (Dumont et al., 2009; Link et al., 2010; Kudapa et al., 2023). Although *GOLS3*, which participates in the synthesis of raffinose, is induced by LT in *A. thaliana* (Taji et al., 2002; Park et al., 2015) as well as its orthologs in our differential expression analysis in *P. sativum* and *C. arietinum*, the equivalent gene has not been identified by the GWAS in *V. faba*. However, our GWAS identified *Vfaba.Hedin2.R1.1g377720*, which orthologs in *P. sativum* and *C. arietinum* upregulated under LT (**Supplementary Material 1**). These genes encode β-amylases and are annotated as part of the maltose biosynthetic process in the chloroplast. When *A. thaliana* is exposed to fluctuations of temperature, the ortholog of these genes, *BAM3*, is induced and correlates with maltose accumulation (Monroe et al., 2014), which improves frost tolerance (Cvetkovic et al., 2021).

Cell wall related genes were enriched among our candidate genes. For example, the ortholog of *Vfaba.Hedin2.R1.1g099840* in *A. thaliana*, *PGAP3B*, encodes a GPI-specific phospholipase that is crucial for the anchoring of proteins to the cell wall/plasma membrane (Bernat-Silvestre et al., 2022). In addition, *Vfaba.Hedin2.R1.5g088200* has for ortholog *ZERZAUST* in *A. thaliana*, a cell wall-localised GPI-anchored β-1,3 glucanase involved in the cell-wall organisation (Vaddepalli et al., 2017). The same protein was highlighted as over-accumulated by shotgun proteomics during cold acclimation of *A. thaliana* (Takahashi et al., 2021). We also revealed *Vfaba.Hedin2.R1.2g079120* as associated with frost tolerance, which encodes a laccase 5/12. The laccase 5/12 genes are predominantly expressed in vascular tissues and in lignified regions of *A. thaliana* (Ehlting et al., 2005; Turlapati et al., 2011). In *P. sativum*, this laccase, and three others (Psat.cameor.v2.2g184650, Psat.cameor.v2.3g95850, Psat.cameor.v2.5g206400), were downregulated in Champagne under LT in this study. It was hypothesized that the downregulation of these genes would result in reduced lignin polymerisation, and increase in cell wall permeability and elasticity contributing to reduced frost damage (Baldwin et al., 2014; Mazurier et al., 2022). The lower expression of a gene encoding a xylan O-acetyltransferase (*Psat.cameor.v2.7g23500*) in Champagne comparatively to Térèse, with an ortholog in *V. faba* (*Vfaba.Hedin2.R1.2g256720*) associated with a frost tolerance MTA, also supports this hypothesis. Interestingly, the ortholog of these genes in *A. thaliana* is *TBL34*, which is involved in the synthesis and deposition of secondary wall cellulose, as is its close paralog, *ESKIMO1* (*ESK1*; Yuan et al., 2016). The *A. thaliana esk1* mutant shows a more compact development than the wild-type and an increased frost tolerance, independently of the canonical *CBF/DREB1* pathway (Xin and Browse, 1998). Finally, increased drought tolerance is observed in *A. thaliana* with low xylan content (Yan et al., 2018). Our GWAS also identified *Vfaba.Hedin2.R1.3g070040*, an ortholog of *MtSEO2* in *M. truncatula* (**Supplementary File 5**; Pélissier et al., 2008). This Faboidae-specific forisome-forming SEO is postulated to play a role in rapidly regulating the flow through the phloem after wounding (Knoblauch et al., 2014). The gene is located on chromosome Vf03, in a syntenic region of chromosome Mt01 of *M. truncatula*. Two of the three orthologs in *P. sativum*, were less expressed under LT in Champagne comparatively to Térèse, while the third gene showed an opposite behaviour. The precise role of these differentially expressed genes in *P. sativum* and whether *V. faba* orthologs have similar functions remain to be elucidated.

In *P. sativum*, some frost-tolerant genotypes, including Champagne, display a delayed floral initiation, conferred by the *Hr* allele of *HIGH RESPONSE TO PHOTOPERIOD* (*HR*; Lejeune-Hénaut et al., 1999) gene, which is an ortholog of *EARLY FLOWERING 3* (*ELF3*) in *A. thaliana* (Weller et al., 2012). The gene in *P. sativum* is located on chromosome Ps05 and has been identified as a major candidate for frost tolerance QTL in Boutet et al. (2023). However, as shown on **Fig. 3**, no syntenic counterparts for these QTL on Ps05 were found on the syntenic chromosome Vf02 in *V. faba*. In fact, in the current version of the *V. faba* genome, the ortholog of *ELF3* is located on a scaffold including only two annotated genes, which has not been integrated in the pseudomolecule. Because the GWAS has been performed using markers successfully positioned on chromosomes only, the possible contribution of *ELF3* to frost tolerance in *V. faba* remains to be studied. Another regulator of flowering time, *SUPPRESSOR OF OVEREXPRESSION OF CONSTANS1* (*SOC1*), which encodes a MADS box transcription factor, has been demonstrated by Seo et al. (2009) to regulate flowering and freezing tolerance in *A. thaliana*. Although its ortholog in *V. faba, Vfaba.Hedin2.R1.5g063160,* was not directly pinpointed by the frost tolerance GWAS, the physically close gene *Vfaba.Hedin2.R1.5g063080* was identified by the three methods, in two environments. The latter displayed no functional annotation and showed no homology with any plant gene. The potential role of *SOC1* in the control of frost tolerance in *V. faba* is therefore worth considering.

The various candidate genes should be the new targets of further investigations and functional validation initiatives as they represent a promising avenue for improvement of frost tolerance in *V. faba* in particular and in cool-season legumes in general.

### Using OrthoLegKB for comparative studies: opportunities, limits and perspectives

The present comparative study of frost tolerance determinism was made possible by the integration of large QTL datasets into the translational database OrthoLegKB. The selection of datasets for integration depended on the availability of markers’ context sequences, which allow reliable alignment and positioning of markers on genomes. The increased popularity of SNP markers in recent years has been instrumental in this process. The use of ontologies has also been crucial, allowing large numbers of datasets to be queried together. The species-neutral TO (Arnaud. et al., 2012; Cooper et al., 2024) was selected to allow the integration of multiple datasets from different cool-season legumes. New terms, including freezing stress response and some more specific related traits, were successfully added to the TO to meet the objectives of this study. Species-specific ontologies were not considered due to the time-consuming manual mapping of traits (Laporte et al., 2016), notably to generalist ontologies. This process is underway (Cooper et al., 2024) and should allow using Crop Ontologies (Arnaud et al., 2016) in the future. We believe that the plant community should take advantage of the potential offered by ontologies. Indeed, while sequencing experiments are increasingly following FAIR guidelines (Stevens et al., 2020), genetic data and experimental protocols are lagging behind, with a variety of formats and some missing information. Some initiatives in this area for human data should provide a model for the plant community (Sollis et al., 2023).

By bringing together heterogeneous data, including functional annotations and RNA-seq experiments, we have also demonstrated that OrthoLegKB can be used to identify robust candidate genes. Including more and, when feasible, high-quality datasets in the database will result in better and more accurate outputs, especially when integrating QTL or expression information from multiple tissues, time points and accessions (Wainberg et al., 2019) and/or species.

The large-scale comparison of freezing tolerance QTL with physiology and phenology QTL located in syntenic regions performed here considered four species, with a minimum of three syntenologous genes under the QTL to form a clique, allowing to go beyond the pairwise synteny implemented in OrthoLegKB. Lowering this threshold to two syntenologs would lead to more noise in the results, partly due to QTL having large confidence intervals and therefore encompassing more genes. Instead, introducing more species with their associated data could lead to finer and more complete resolution. This step is underway with an increasing number of species being integrated into OrthoLegKB. The implementation of new modules to handle co-expression networks and polymorphism information is also under serious consideration. Such data types, if integrated into OrthoLegKB, would bring more power to functionally connect genes and highlight putative causal genetic variations (Li et al., 2023). This study paves the way for using OrthoLegKB to other stresses and/or morphological and physiological traits and certainly for using Ortho_KB to create new instances allowing to study other groups of plant species.

## Author contributions

BI managed data annotation, integration and analysis, and drafted the manuscript. JK performed the GWAS analysis. IL-H, GB and GA contributed to data collection and annotation. PM supervised field work and J-BM-R managed the plant material and performed phenotyping. NT contributed to the scientific management of this work and was involved in drafting and writing the manuscript. JB contributed to the interpretation of the results and managed funding acquisition. All authors have read and approved the submitted version.

## Acknowledgements

The authors are grateful to Planteome and in particular to Laurel Cooper for her help in integrating new ontology terms and to the INRAE URGI team and in particular to Raphaël-Gauthier Flores for his help in updating OrthoLegKB. They are also thankful to the INRAE experimental units based in Dijon and Theix and to the Agri Obtentions team in Orsonville for carrying out the field experiments and contributing to the collection of phenotyping data.

---

**Supplementary Figure 1.**
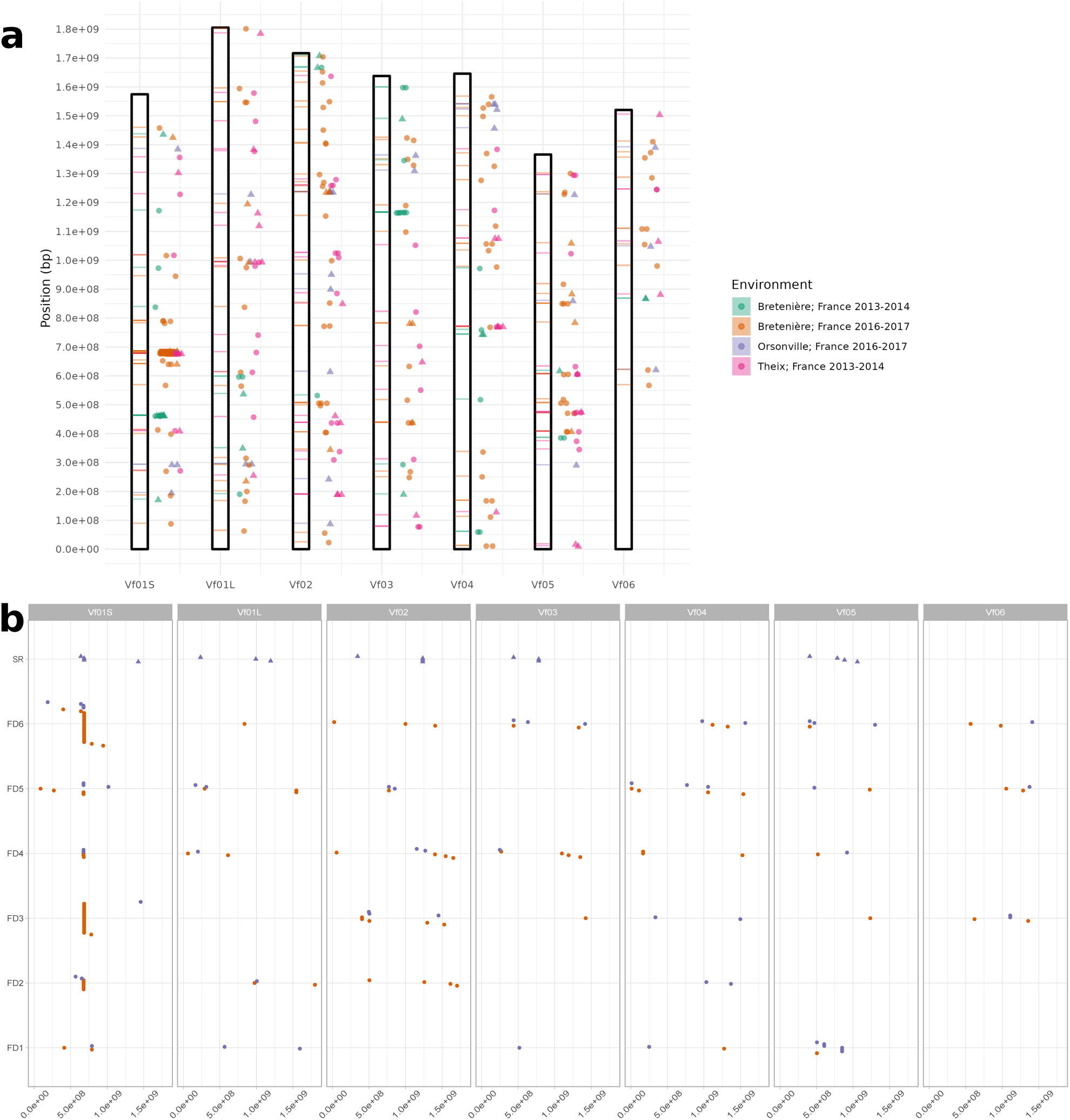
Genomic positions of SNP markers significantly associated with frost tolerance on *V. faba* chromosomes. a) Marker-trait associations (MTAs) revealed using data collected in four environments. Significant markers are represented by both horizontal segments and shapes on the right side of the chromosome. MTAs for frost damage and survival rate traits are represented by dots and triangles, respectively. The colours of the shapes and of the horizontal segments indicate the environment in which the association was detected. b) MTAs relative to phenotyping data collected in Bretenière 2016-2017. The colours of the shapes indicate the GWAS method that allowed the detection of the MTA.

**Supplementary Figure 2.**
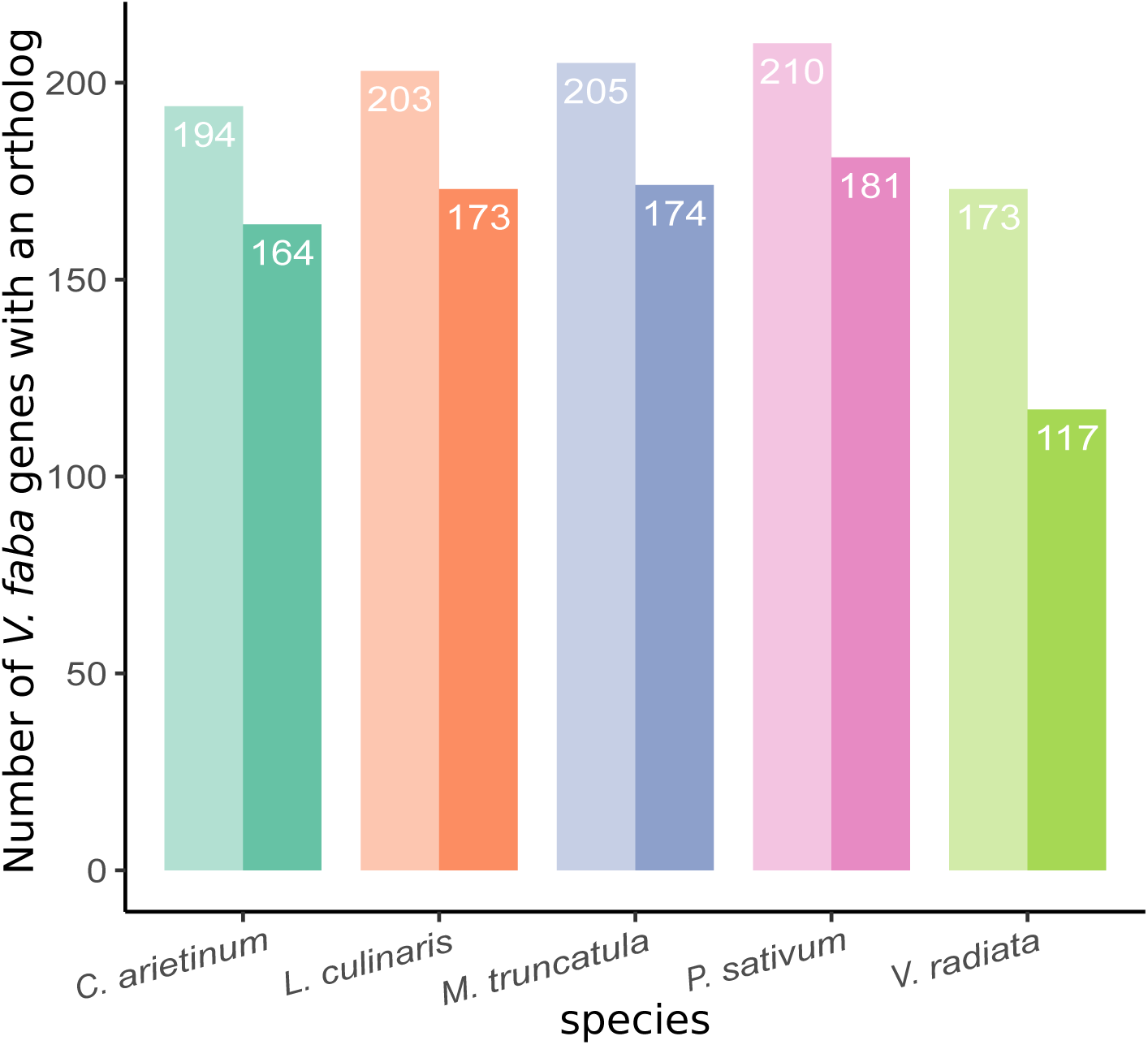
Conservation of gene content between *Vicia faba* and other sister legume species in genomic regions containing QTL controlling frost tolerance. Orthologs for *V. faba* genes harbouring markers significantly associated with frost tolerance were searched in *Cicer arietinum*, *Lens culinaris*, *Medicago truncatula*, *Pisum sativum* and *Vigna radiata* using OrthoLegKB. The total number of *V. faba* genes that have at least one gene part of the same orthogroup in the sister species, is shown in each case on the lighter bars. The total number of *V. faba* genes with syntenic orthologs is shown in darker shades.

**Supplementary Figure 3.**
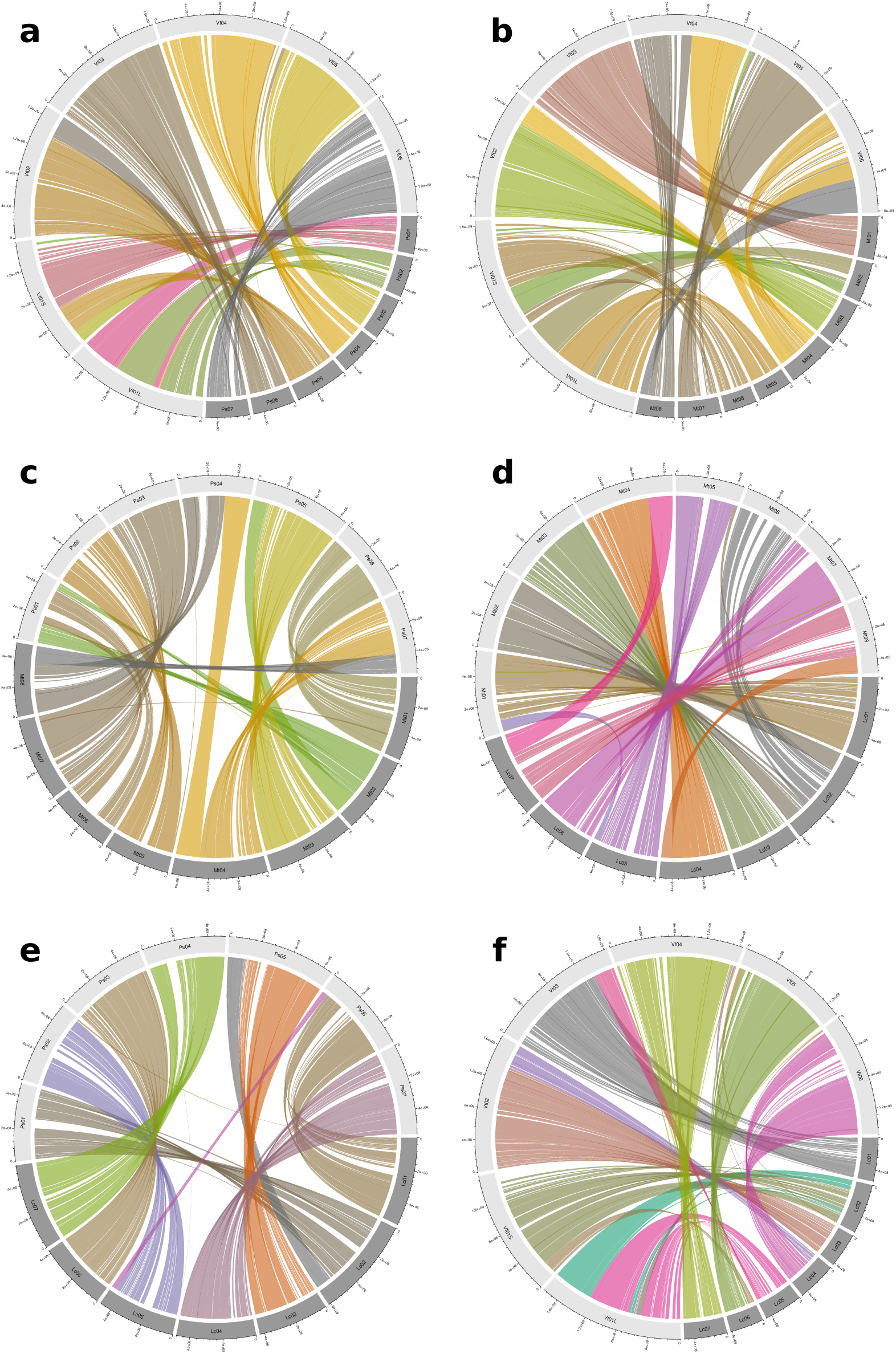
Synteny between legume species relatively close to *Vicia faba*. Synteny is represented using Circos plots. Circos plots are between *V. faba* and *Pisum sativum* (a), *V. faba* and *Medicago truncatula* (b), *M. truncatula* and *P. sativum* (c), *V. faba* and *Lens culinaris* (d), *L. culinaris* and *P. sativum* (e), and *L. culinaris and M. truncatula* (f). Each Circos plot shows the chromosomes on the outer track. The innermost links indicate syntenic genomic blocks. The total length of *M. truncatula* chromosomes has been increased by a factor of 10 for readability.

**Supplementary Figure 4.**
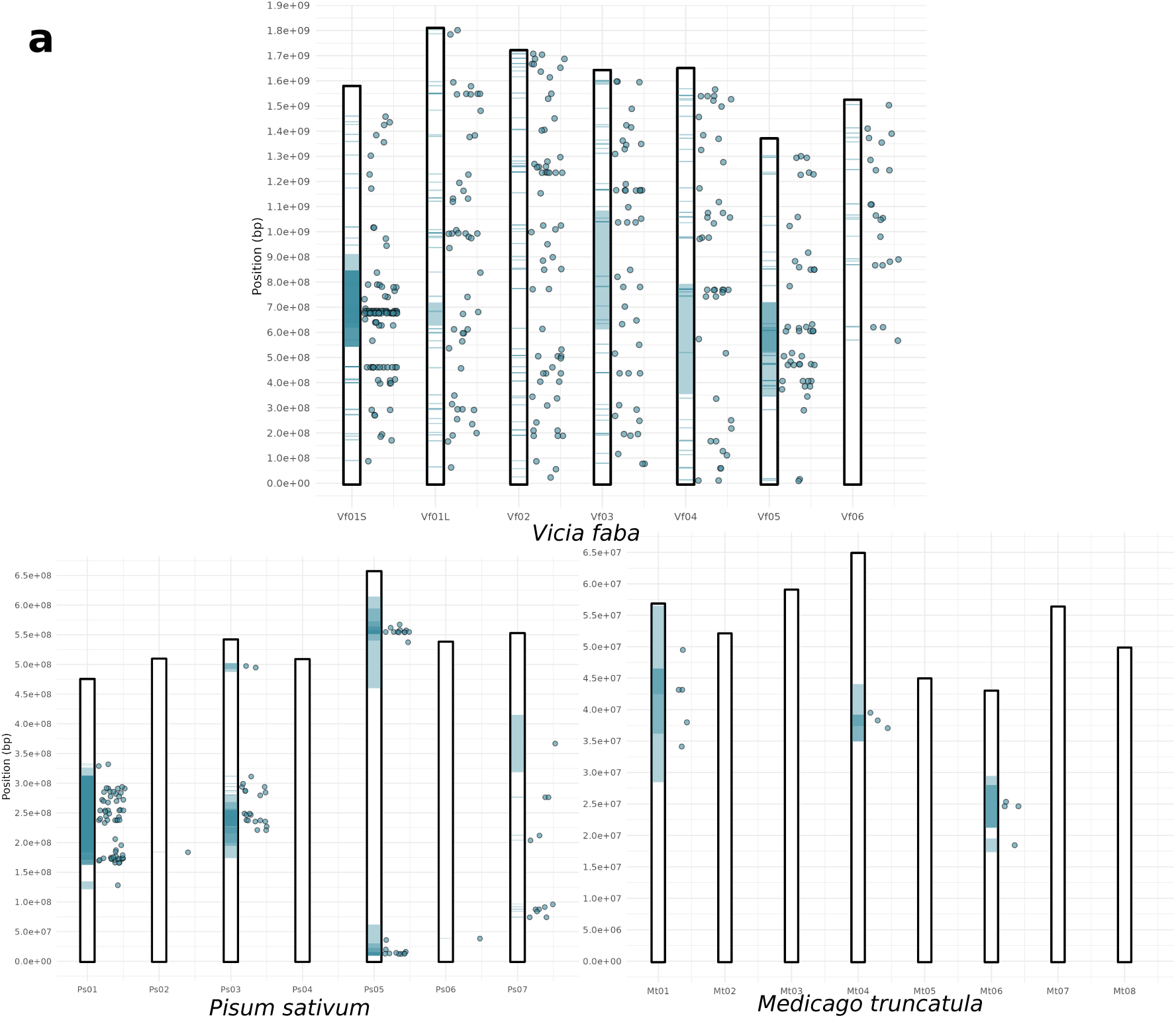
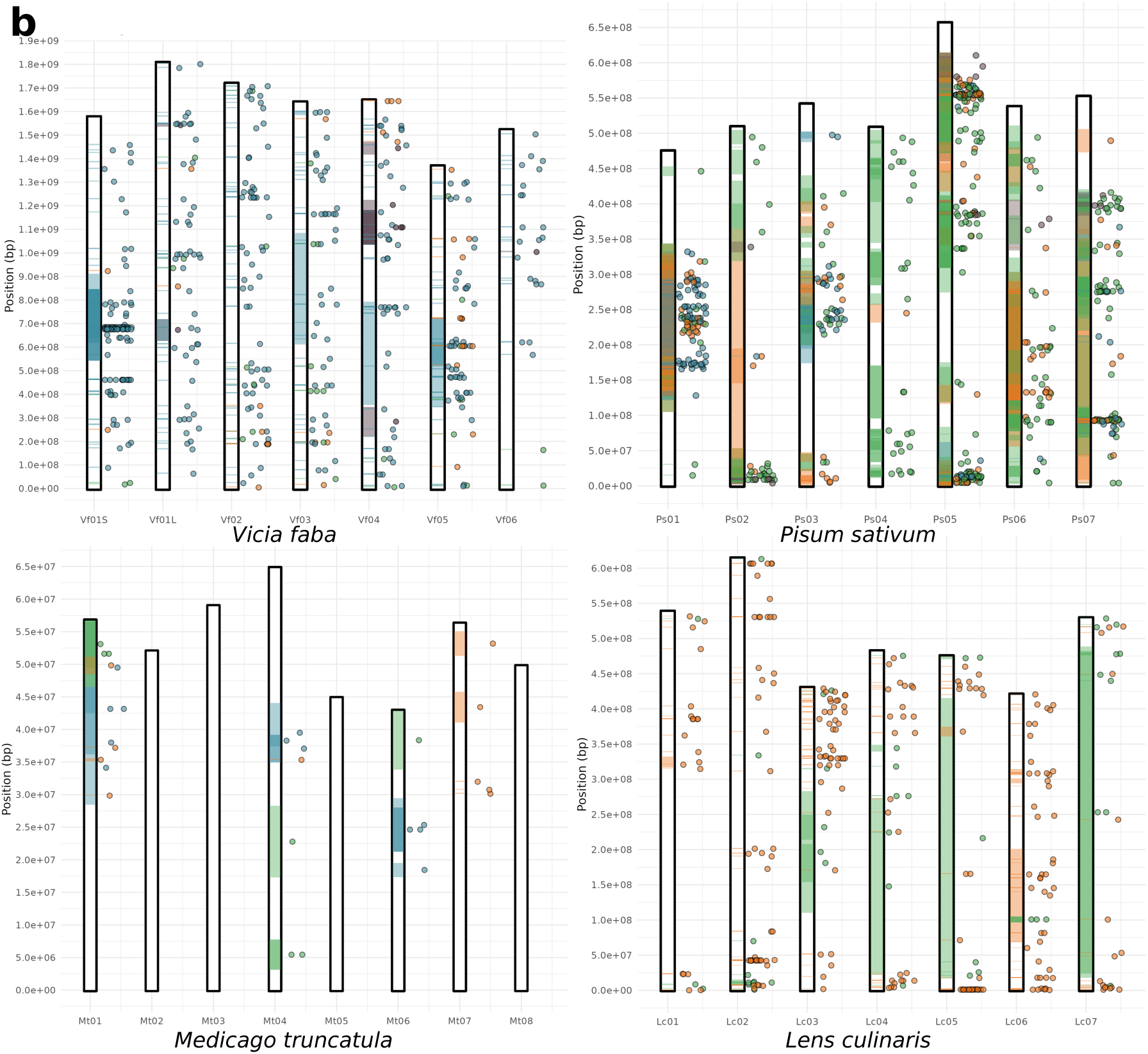
Overview of QTL used in this study, on chromosomes of the respective legume species. a) QTL for frost tolerance (light blue) on the chromosomes of *Vicia faba*, *Pisum sativum* and *Medicago truncatula*. b) QTL on the chromosomes of *V. faba*, *P. sativum*, *M. truncatula* and *Lens culinaris*. QTL for frost tolerance are shown in light blue, while QTL for flowering time, fat and carbohydrate content, and morphological traits are shown in orange, taupe, and green, respectively. Chromosomes are represented by vertical bars with their respective names below. Each QTL is represented by both a rectangle indicating its confidence interval on the chromosome and by a dot to the right side of the chromosome indicating the peak position. The colours of the rectangles and of the dots refer to the respective traits. To make the GWAS peaks visible in this figure, their intervals were extended to 1,000,000, 5,000,000, and 150,000 bp for *P. sativum, V. faba, and M. truncatula*, respectively.

**Supplementary Figure 5.**
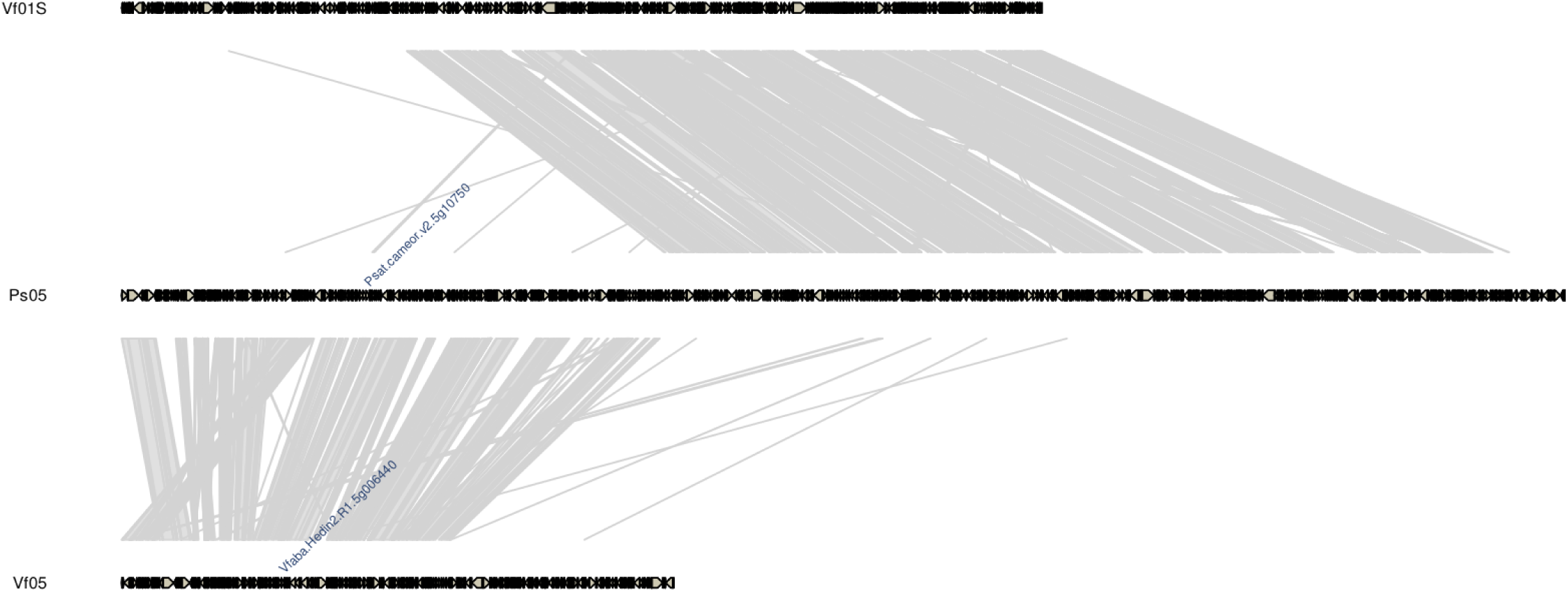
Microsynteny between the genomic region containing the *Le* locus on *Pisum sativum* chromosome Ps05 and *Vicia faba* chromosomes Vf01S and Vf05. The gene controlling internode length in *P. sativum*, designated *Le*, is *Psat.cameor.v2.5g10750*. *Vfaba.Hedin2.R1.5g006440* is the *V. faba* syntenic ortholog of *Psat.cameor.v2.5g10750*. Since the two species have large genome sizes and more or less large intergenic distances, the intergenic regions are not shown in this figure. However, the gene sizes remain proportional.

