## Supplementary Material 1 for "Genome-wide association study of frost tolerance in *Vicia faba* reveals syntenic loci in cool-season legumes and highlights relevant candidate genes"

**Supplementary Material 1**: Differential expression of *Pisum sativum* and *Cicer arietinum* genes under low temperature treatment in the datasets of Bahrman et al. (2019) and Akbari et al. (2023). The comparison was performed for each species between a tolerant genotype and a susceptible genotype.

### Material and methods

#### Functional annotation enrichment analysis

Functional annotations including MapMan annotation (Thimm et al., 2004; Schwacke et al., 2019), Gene Ontology (GO) annotation (Ashburner, 2000; The Gene Ontology Consortium et al., 2023), and InterPro annotation (Jones et al., 2014; Blum et al., 2021) were performed for *Pisum sativum*, *Vicia faba,* and *Lens culinaris* genes as described in Imbert et al. (2023). Annotations were queried in OrthoLegKB for each gene using its unique gene ID. The enrichment of gene functions at specific loci or in expression data was evaluated by conducting gene enrichment analyses based on GO terms in R v4.3.3 (R Core Team, 2024) with topGO v2.54.0 (Alexa and Rahnenfuhrer, 2023), using Fisher's test and the "classic" algorithm of the package for the handling of the GO graph structure (The Gene Ontology Consortium et al., 2023). The results were then visualized using the R package clusterProfiler v4.10.0 (Wu et al., 2021). For MapMan bins (Schwacke et al., 2019), the enrichment was done using the enrichplot package v1.22.0 (Guangchuang Yu, 2018).

#### Differential gene expression analysis

Twenty-four *P. sativum* RNA-seq samples (NCBI BioProject PRJNA543764), corresponding to 2 genotypes × 2 treatments × 3 sampling times × 2 biological replicates, from Bahrman et al. (2019) were used in this study. Briefly, Bahrman et al. (2019) considered a frost-tolerant (Champagne) and a frost-susceptible (Térèse) accession and grew them for a nursery period of 21 days under control conditions (N; 20°C day/14°C night at 500 $\mu mol.m^{-2}.s^{-1}$ photosynthetic photon flux with a 10 h photoperiod). Two groups were then defined, with one remaining in N conditions while the other was exposed to low temperature (LT; 8°C day/2°C night at 250 $\mu mol.m^{-2}s^{-1}$ photosynthetic photon flux with a 10 h photoperiod). Aerial parts of the plants were sampled at equivalent developmental stages of both treatments, i.e. after 20 (T0), 22 (T1) and 27 days (T2) in the N treatment and after 20 (T0), 23 (T1) and 36 (T2) in the LT treatment. Raw pseudo-counts for each RNA-seq sample were generated using Salmon (Patro et al., 2017) implemented in the nf-core/rnaseq pipeline v.3.12.0 (Patel et al., 2023), converted to integers, and then subjected to differential expression analysis using the DESeq2 R package v1.42.1 (Love et al., 2014). Each sample was annotated using Planteome Plant Ontology (PO) and Plant Experimental Conditions ontology (PECO; Cooper et al. 2024) and the pseudo-counts for each gene integrated in OrthoLegKB. Data corresponding to T1 and T2 sampling dates were submitted to multifactorial analysis using DESeq2 in order to highlight (1) differentially expressed genes (DEGs) under low non negative temperatures within each genetic background considered and (2) DEGs under LT or N when comparing Champagne and Térèse genotypes. DEGs with an adjusted $p$-value < 0.05 and a | log2 fold change | > 1 were retained for further analysis.

The same procedure as described above was used to analyse *C. arietinum* RNA-seq samples from

Akbari et al. (2023), obtained from the frost-tolerant genotype Saral and frost-susceptible genotype ILC533 (available in NCBI BioProjects PRJNA905065 and PRJNA903665). The queries to retrieve transcriptomic data from OrthoLegKB are available in **Supplementary File 2**.

### Results

#### Identification of differentially expressed genes in response to low temperature treatment in *Pisum sativum* and colocalisation of their orthologous genes with frost tolerance-related QTL in *Vicia faba*

##### Reanalysis of a transcriptomic dataset on gene expression response to low temperature in *P. sativum*

In order to find potential candidate genes for the frost tolerance marker-trait associations (MTAs) identified by GWAS in *V. faba*, and given the lack of an adequate transcriptomic dataset for this species, we decided to use the available RNA-seq resources obtained in *P. sativum* under LT treatment and attempt a knowledge transfer to *V. faba*, which is easily enabled by OrthoLegKB. Indeed, Bahrman et al. (2019) published, before the release of the first genome assembly of *P. sativum*, an RNA-seq dataset derived from the aerial parts of young *P. sativum* plants and identified molecular determinants for the response to LT in two accessions contrasted for frost tolerance. Champagne is known to be frost-tolerant while Térèse is frost-susceptible (Bahrman et al., 2019).

As listed in **Supplementary Table 5** and depicted in **Supplementary Material Fig. 1a**, our transcriptomic reanalysis using the latest assembly of *P. sativum* cv. Caméor revealed 8,252 differentially expressed genes (DEGs) either between genotypes or treatments for the same genotype. At the sampling time T1, we revealed 1,657 and 1,665 DEGs for Térèse and Champagne respectively, when submitted to LT, with more upregulated genes in Champagne than in Térèse. When comparing the differential expression of both genotypes at T1, we showed that 364 downregulated and 437 upregulated genes were shared between them. At the sampling point T2, the number of downregulated genes increased by factor of 1.6 or more in both genotypes, while the number of upregulated genes slightly decreased. Among the downregulated genes, 447 were specific to T2 and shared between both genotypes, while 468 were specific to Champagne. As for the upregulated genes, approximately the same number of DEGs specific to this time point and to each genotype were found. The two sampling points were characterized by strong differences in the DEGs, few being shared at the intra-genotype scale. When comparing gene expression across genotypes, we showed that 1,141 DEGs specific to T2 were upregulated in Térèse, and 703 DEGs specific to T2 were upregulated in Champagne. Only 111 genes were upregulated in Champagne compared to Térèse under LT at both sampling points (**Supplementary Material Fig. 1b**). To identify the main biological processes involved in the response to LT and better understand the difference of response between contrasted genotypes, we performed a GO and MapMan enrichment analysis for DEGs. When comparing Champagne to Térèse, we found at both T1 and T2 a statistical significance for several GO terms involved in the response to biotic and abiotic stimuli, such as cold, and light intensity, with the involvement of the karrikin (strigolactone-like) hormone. The

MapMan bins enrichment revealed genes encoding glycosyltransferase, class tau glutathione S-transferase, DUF26 protein kinase and leucine-rich repeat immune receptors (NLRs) and many oxidoreductase enzymes such as a leucine-rich receptor-like protein kinase (**Supplementary Material Fig. 2a and b**). Some GO enrichments were specific to T2, including for the response to lipid, abscisic acid and alcohol. MapMan bins were further enriched for genes encoding CBF/DREB1 transcription factors, in Champagne compared to Térèse (**Supplementary Material Fig. 2b**). This enrichment of *CBF/DREB1* genes was already observed in Champagne at T1 compared to its own control (data not shown). To specifically observe the enriched functions of genes induced by LT at T2 where more genes annotated by the GO term "response to cold" were detected, we decided to subtract genes that were already differentially expressed between the studied accessions under control conditions. This allowed to also identify that cell wall remodelling process seems to play a role in adaptive cold acclimation (**Supplementary Material Fig. 3**). Indeed, we observed a significative enrichment of terms "polysaccharide metabolic process", "xylan biosynthetic process", "plant-type secondary cell wall biogenesis".

#### Highlighting low temperature-responsive *Pisum sativum* genes having orthologs underlying frost tolerance QTL in *Vicia faba*

To shade light on candidate genes underlying *V. faba* frost tolerance associated QTL, we filtered *P. sativum* genes differentially expressed either in response to LT or between contrasted genotypes for their response to LT and kept only those having at least one ortholog in the vicinity of the frost tolerance-associated MTAs in *V. faba*. In total, 141 *P. sativum* genes, hereafter referred to as qDEGs, matching these criteria were selected (**Supplementary Material Fig. 4a**). As depicted in **Supplementary Material Fig. 4b**, for intra-genotype comparisons, we found 60 downregulated qDEGs, with 19 specific to the T1 sampling time, and 34 specific to T2. Regarding upregulated genes, we identified 36 qDEGs, with eight specific to T1 and 12 to T2. While most downregulated qDEGs were specific to each sampling time and genotype, more than 18% of upregulated qDEGs were shared across sampling times and genotypes. The comparison of genotypes informed that LT induced the expression of 55 genes with a higher expression in Térèse compared to Champagne, and 53 genes with a higher expression in Champagne compared to Térèse (**Supplementary Material Fig. 4b**). All commonalities displayed on the upset plots in **Supplementary Material Fig. 4** and underlying genes are described in **Supplementary Material Table 1**.

A functional enrichment with MapMan allowed to highlight some common functions of qDEGs compared to all genes of the *P. sativum* genome (**Supplementary Fig. 5**). The enriched functions showed that qDEGs included five upregulated genes annotated as *CBF/DREB1*, four genes annotated as *WRKY*, with three of them downregulated only in Térèse under LT at T2. We also have detected seven genes encoding EC 3-2 glycosylases. These genes seemed to be more expressed in Champagne in control conditions and to exhibit a lower expression in Térèse at T2. We also have identified two genes encoding auxin efflux transporters, with one more expressed in Champagne than in Térèse in control conditions, but downregulated under LT at T2 in Champagne. We detected three consecutive genes on the genome, all upregulated in at least one LT contrast, annotated as class tau glutathione S-transferase. The enrichment of EC 1-13 oxidoreductases was supported by three *P. sativum* genes that were strongly downregulated in Champagne and Térèse under LT at T2 and also showed a lower expression in Champagne compared to Térèse at the same sampling time.

Although "xylan O-acetyltransferase" was not enriched for qDEGs, we identified one gene encoding for this enzyme which showed a higher expression in Térèse compared to Champagne at T1 under LT and at T2 in both conditions.

Using the same methodology, we searched for qDEG genes in the RNA-seq dataset from Akbari et al. (2023) in *C. arietinum*, comparing Saral, a frost-tolerant genotype and ILC533, a frost-susceptible genotype (**Supplementary Table 6**). Crossing the results from *P. sativum* and *C. arietinum* led to a list of 16 genes with qDEGs up- or downregulated in the same direction in both tolerant accession or both susceptible accession (**Supplementary Table 11**). As in the paper of Akbari et al. (2023), we found a WRKY encoding gene, *Ca_v2.0_04345*, ortholog of *Vfaba.Hedin2.R1.1g229680*, which was downregulated under LT in both susceptible genotypes. We also have identified genes encoding for enzymes, namely *Vfaba.Hedin2.R1.1g18380* encoding a class tau glutathione S-transferase, *Vfaba.Hedin2.R1.2g079120* encoding a laccase multi-cooper oxidoreductase (LAC5/LAC12), *Vfaba.Hedin2.R1.1g377720* encoding a β-amylase, *Vfaba.Hedin2.R1.5g086480* encoding an EC 2-4 glycosyltransferase, *Vfaba.Hedin2.R1.6g119600* encoding an EC_1-14 oxidoreductase, *Vfaba.Hedin2.R1.2g266920* encoding an acyl-coA desaturase (ADS/FAD5) and *Vfaba.Hedin2.R1.2g264600* with no known function. Interestingly, we also have detected a gene encoding a sieve element occlusion protein (SEO), with orthologous genes showing a higher expression in the tolerant genotypes under LT compared to susceptible genotypes and for which the *V. faba* ortholog, *Vfaba.Hedin2.R1.3g070040*, was bearing three MTAs for frost damage and winter survival identified in Breteniëre 2016-2017. Looking at the previously described 19 regions dense in MTAs in *V. faba* (**Supplementary Table 2** and **Fig. 2**) revealed only one gene with qDEGs in *P. sativum* and *C. arietinum*, *Vfaba.Hedin2.R1.5g073960*, encoding an ER luminal lectin chaperone, which was supported by MLM only. Several *V. faba* genes in MTA-rich regions were supported by qDEGs in *P. sativum* only, including the *CBF/DREB1* genes *Vfaba.Hedin2.R1.1g099680*, *Vfaba.Hedin2.R1.1g099880*, *Vfaba.Hedin2.R1.1g100240* and *Vfaba.Hedin2.R1.1g101280*. Other genes such as *Vfaba.Hedin2.R1.2g085600* encoding a peptidase, *Vfaba.Hedin2.R1.4g116520* encoding a phosphoinositide transfer protein, *Vfaba.Hedin2.R1.4g116800* encoding an EC-1-14 oxidoreductase, *Vfaba.Hedin2.R1.4g116920* encoding a kinetochore SPC25-related protein, *Vfaba.Hedin2.R1.5g073520* encoding an EC-3-4 hydrolase and *Vfaba.Hedin2.R1.5g073960* encoding an ER luminal lectin chaperone were also found.

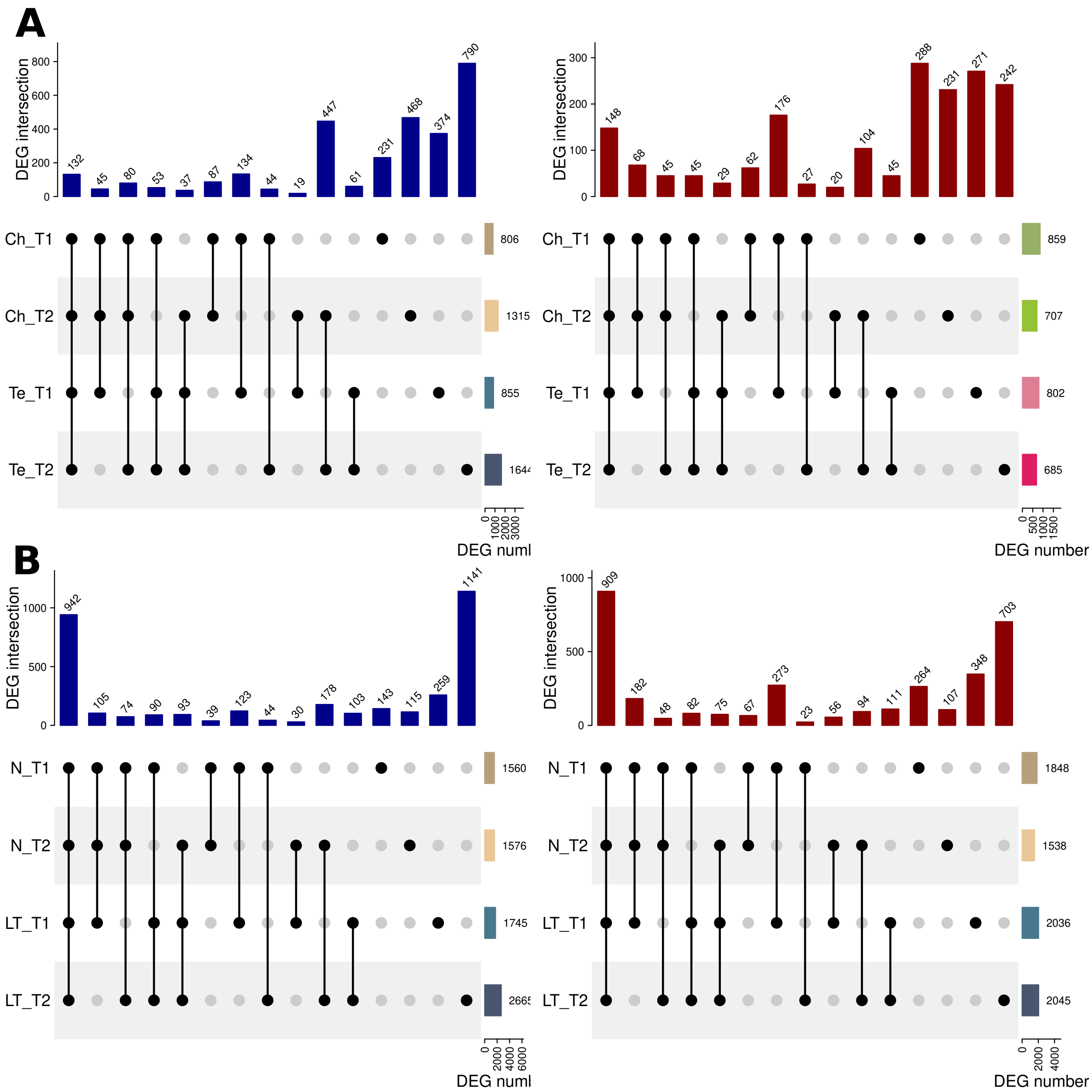


**Supplementary Material Figure 1**. Upset plots of differentially expressed genes in *Pisum sativum* accessions Champagne (Ch) and Térèse (Te) at two experimental time points, T1 and T2, following exposure (LT) or not (N) to low temperature treatment. An unconnected black dot indicates differentially expressed genes found only in this condition, while connected dots indicate the intersection of the gene sets expressed in two conditions or more. The bar graph on the right side of each upset plot shows the total number of differentially expressed genes in each condition. a) Upregulated (left panel, blue) and downregulated (right panel, red) genes in Champagne and in Térèse under low temperatures compared to the respective control. b) Upregulated (left panel, blue) and downregulated (right panel, red) differentially expressed genes in Champagne compared to Térèse at equivalent control and sampling times under low temperatures.

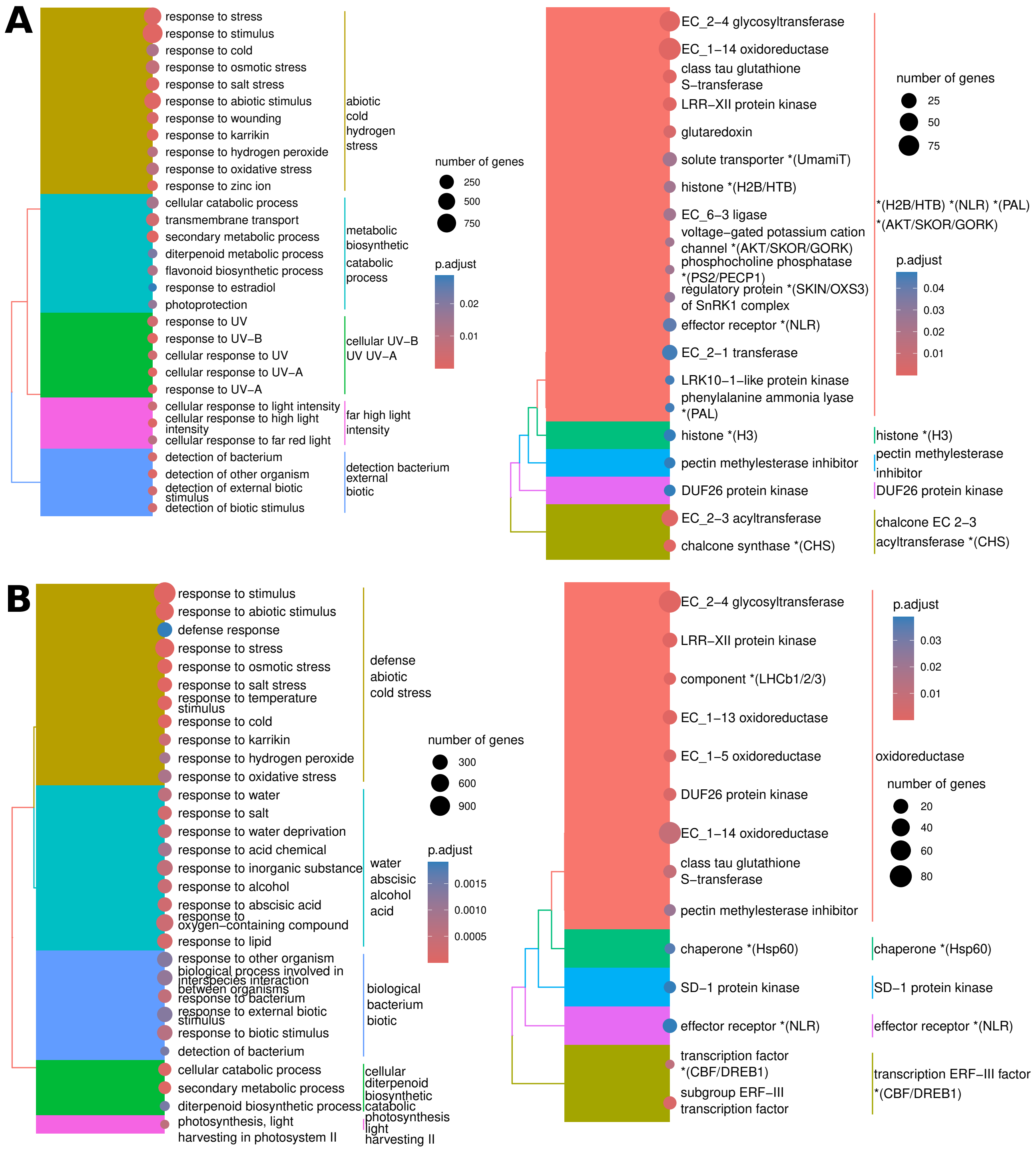


**Supplementary Material Figure 2**. Functional annotation enrichment of differentially expressed genes in *Pisum sativum* accessions Champagne compared to Térèse upon low temperature exposures at two time points, T1 (a) and T2 (b). The left panels were constructed using enriched GO terms, while the right panels were constructed using enriched MapMan bins. The size of the circle of each node is relative to the number of genes carrying the annotation, and the colour of the circle reflects the adjusted p-value after false discovery rate correction.

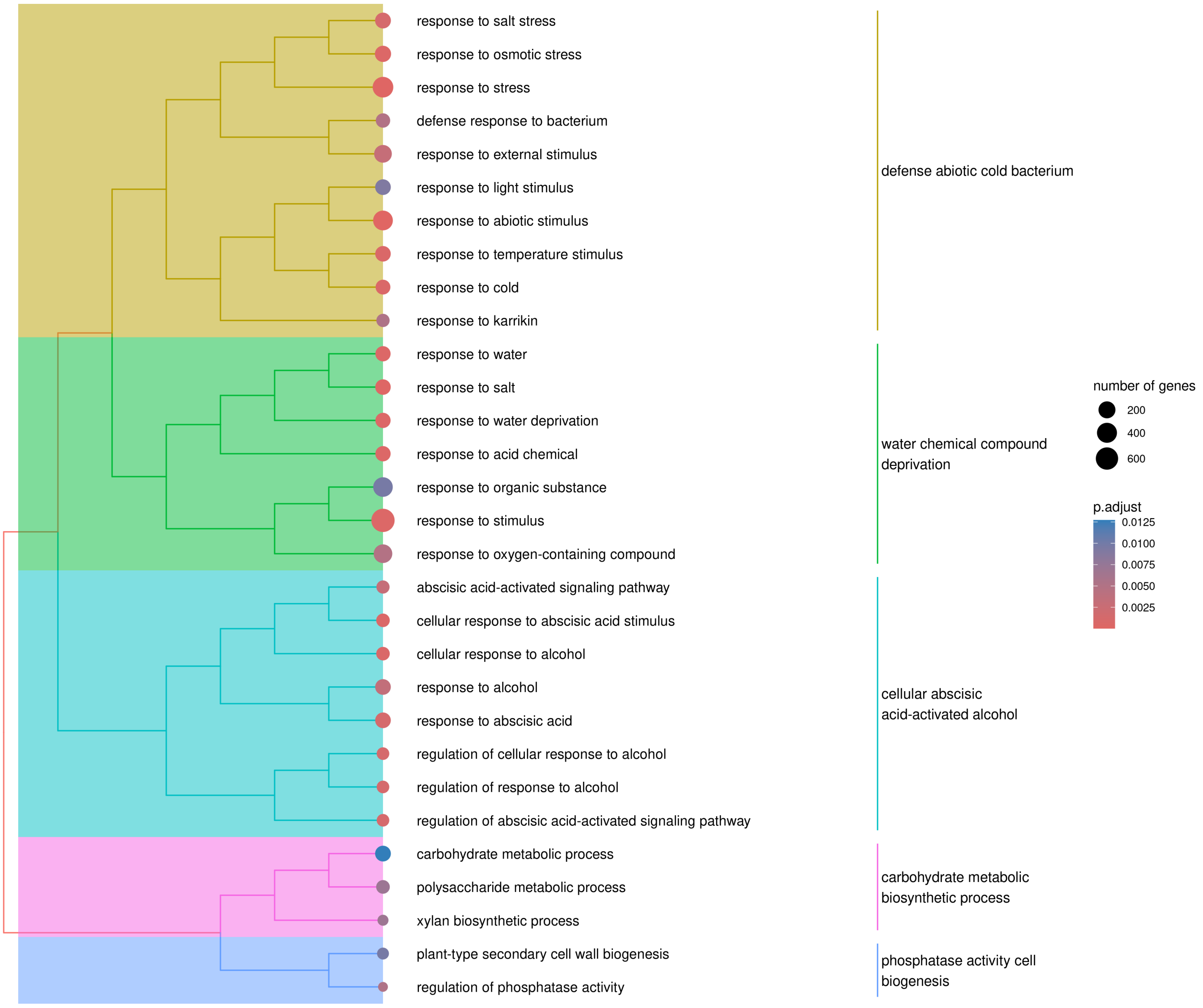


**Supplementary Material Figure 3**. Hierarchical clustering analysis of Gene Ontology terms enrichment revealed by analysis of significantly up- or downregulated genes in the *Pisum sativum* frost-tolerant accession Champagne compared to the frost-susceptible accession Térèse after exposure to low temperature at experimental time point T2. Genes differentially expressed between the two genotypes in control conditions were not taken into account (see Materials and Methods). The size of the dots depends on the number of genes carrying the annotation, and the colour of the dots reflects the adjusted p-value after false discovery rate correction.

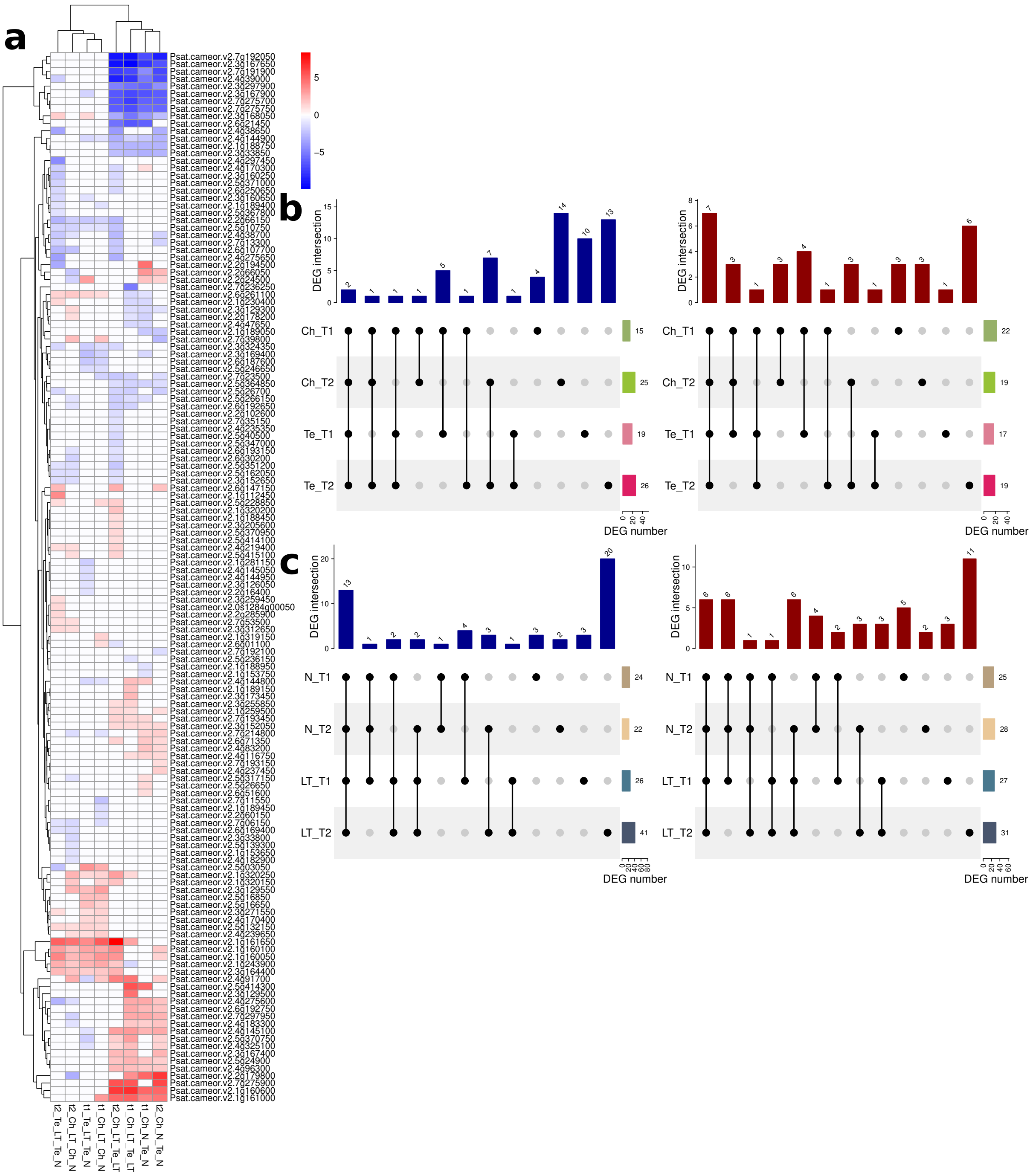


**Supplementary Material Figure 4**. Upset plots of differentially expressed genes in *Pisum sativum* with an ortholog in *V. faba* harbouring a marker-trait association from the GWAS analysis for frost tolerance. The *P. sativum* accessions Champagne (Ch) and Térèse (Te) have been sampled at two experimental time points, T1 and T2, following exposure (LT) or not (N) to low temperature treatment. An unconnected black dot indicates differentially expressed genes found only in this condition, while connected dots indicate the intersection of the gene sets expressed in two conditions or more. The bar graph on the right side of each upset plot shows the total number of differentially expressed genes in each condition. a) Heatmap of the base 2 log fold-change of differentially expressed genes. b) Upregulated (left panel, blue) and downregulated (right panel, red) genes in Champagne and in Térèse under low temperatures compared to the respective control, c) Upregulated (left panel, blue) and downregulated (right panel, red) differentially expressed genes in Champagne compared to Térèse at equivalent control and sampling times under low temperatures.

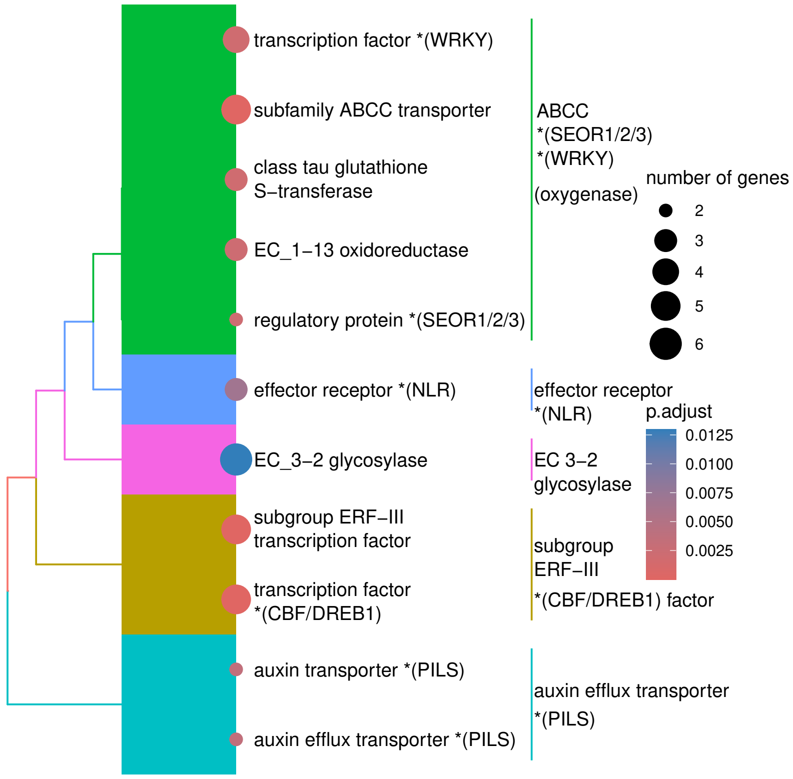


**Supplementary Material Figure 5**. Hierarchical clustering analysis of MapMan bin enrichment using the 95 *Pisum. sativum* genes with an ortholog in *Vicia faba* harbouring a marker-trait association from the GWAS analysis for frost tolerance. The size of the dots depends on the number of genes carrying the annotation, and the colour of the dots reflects the adjusted p-value after false discovery rate correction.

Akbari, A., Ismaili, A., Amirbakhtiar, N., Pouresmael, M., and Shobbar, Z.-S. (2023). Genome-wide transcriptional profiling provides clues to molecular mechanisms underlying cold tolerance in chickpea. *Sci Rep* 13, 6279. doi: 10.1038/s41598-023-33398-3

Alexa, A., and Rahnenfuhrer, J. (2023). topGO. *Bioconductor*. Available at: http://bioconductor.org/packages/topGO/ (Accessed April 24, 2024).

Ashburner (2000). Gene ontology: tool for the unification of biology. *Nat Genet*. Available at: https://www.ncbi.nlm.nih.gov/pmc/articles/PMC3037419/

Bahrman, N., Hascoët, E., Jaminon, O., Dépta, F., Hû, J.-F., Bouchez, O., et al. (2019). Identification of Genes Differentially Expressed in Response to Cold in Pisum sativum Using RNA Sequencing Analyses. *Plants* 8, 288. doi: 10.3390/plants8080288

Blum, M., Chang, H.-Y., Chuguransky, S., Grego, T., Kandasaamy, S., Mitchell, A., et al. (2021). The InterPro protein families and domains database: 20 years on. *Nucleic Acids Research* 49, D344–D354. doi: 10.1093/nar/gkaa977

Cooper, L., Elser, J., Laporte, M.-A., Arnaud, E., and Jaiswal, P. (2024). Planteome 2024 Update: Reference Ontologies and Knowledgebase for Plant Biology. *Nucleic Acids Research* 52, D1548–D1555. doi: 10.1093/nar/gkad1028

Guangchuang Yu (2018). enrichplot. doi: 10.18129/B9.BIOC.ENRICHPLOT

Imbert, B., Kreplak, J., Flores, R.-G., Aubert, G., Burstin, J., and Tayeh, N. (2023). Development of a knowledge graph framework to ease and empower translational approaches in plant research: a use-case on grain legumes. *Front. Artif. Intell.* 6. doi: 10.3389/frai.2023.1191122

Jones, P., Binns, D., Chang, H.-Y., Fraser, M., Li, W., McAnulla, C., et al. (2014). InterProScan 5: genome-scale protein function classification. *Bioinformatics* 30, 1236–1240. doi: 10.1093/bioinformatics/btu031

Love, M. I., Huber, W., and Anders, S. (2014). Moderated estimation of fold change and dispersion for RNA-seq data with DESeq2. *Genome Biol* 15, 550. doi: 10.1186/s13059-014-0550-8

Patel, H., Ewels, P., Peltzer, A., Botvinnik, O., Sturm, G., Moreno, D., et al. (2023). nf-core/rnaseq: nf-core/rnaseq v3.10.1 - Plastered Rhodium Rudolph. doi: 10.5281/zenodo.7505987

Patro, R., Duggal, G., Love, M. I., Irizarry, R. A., and Kingsford, C. (2017). Salmon provides fast and bias-aware quantification of transcript expression. *Nat Methods* 14, 417–419. doi: 10.1038/nmeth.4197

R Core Team (2024). R: A Language and Environment for Statistical Computing. Available at: https://www.R-project.org/

Schwacke, R., Ponce-Soto, G. Y., Krause, K., Bolger, A. M., Arsova, B., Hallab, A., et al. (2019). MapMan4: A Refined Protein Classification and Annotation Framework Applicable to Multi-Omics Data Analysis. *Molecular Plant* 12, 879–892. doi: 10.1016/j.molp.2019.01.003

The Gene Ontology Consortium, Aleksander, S. A., Balhoff, J., Carbon, S., Cherry, J. M., Drabkin, H. J., et al. (2023). The Gene Ontology knowledgebase in 2023. *Genetics* 224, iyad031. doi: 10.1093/genetics/iyad031

Thimm, O., Bläsing, O., Gibon, Y., Nagel, A., Meyer, S., Krüger, P., et al. (2004). mapman: a user-driven tool to display genomics data sets onto diagrams of metabolic pathways and other biological processes. *The Plant Journal* 37, 914–939. doi: 10.1111/j.1365-313X.2004.02016.x

Wu, T., Hu, E., Xu, S., Chen, M., Guo, P., Dai, Z., et al. (2021). clusterProfiler 4.0: A universal enrichment tool for interpreting omics data. *The Innovation* 2, 100141. doi: 10.1016/j.xinn.2021.100141
