## Supplementary Files for "Genome-wide association study of frost tolerance in *Vicia faba* reveals syntenic loci in cool-season legumes and highlights relevant candidate genes"

**Supplementary File 1. Cypher query used in OrthoLegKB to access information from the current GWAS presented as Supplementary Table 1**

```
MATCH (r:Resource)<-[:HAS_TRAIT]-(q:QTL)-[pos]->(c:Chromosome),
(g:Gene)-->(q)-->(d:Dataset)
WHERE d.dataset_id = 'faba_frost_gwas2023'
OPTIONAL MATCH (s:Site)<--(q)-->(y:Year)
OPTIONAL MATCH (g)-->(:RNA)-->(m:Resource:MapMan)
OPTIONAL MATCH (g)-->(:RNA)-->(go:Resource:GO)
OPTIONAL MATCH (g)-->(:RNA)-->(p:Protein)-->(f:FunctionalAnnotation)
WHERE f.source in ['Pfam', 'PANTHER', 'SUPERFAMILY']
RETURN DISTINCT c.chromosome_id as chromosome, q.qtl_id as qtl_id, q.peakmarker_id as
SNP, q.model as model, q.pvalue as pvalue, pos.start as start, pos.end as end, s.site_id as site,
y.year_id as year, r.label as trait, g.gene_id as gene, g.start as gene_start, g.end as gene_end,
COLLECT(DISTINCT m.label) as mapman, COLLECT(DISTINCT go.label) as go,
COLLECT(DISTINCT f.description) as interpro ORDER BY chromosome, start
```

**Supplementary File 2. Cypher queries used in OrthoLegKB to retrieve RNA-seq data and metadata**

```
MATCH (d:Dataset:RNASeq)<--(c:Condition:RNASeq)<--(s:Sample:RNASeq)
WHERE d.dataset_id = 'PRJNA543764'
OPTIONAL MATCH (s)<-[expr]-(g:Gene)
WHERE g.genome = 'psat'
RETURN DISTINCT g.gene_id as gene,
s.sample_id as sample_id, expr.count as count
```

```
MATCH (d:Dataset:RNASeq)<--(c:Condition:RNASeq)<--(s:Sample:RNASeq),
(c)-[rel]->(r:Resource)
WHERE d.dataset_id = 'PRJNA543764'
RETURN DISTINCT s.sample_id as sample_id, c.condition_id as condition_id,
r.label as condition_label, type(rel) as condition_type
```

**Supplementary File 3. Cypher queries used in OrthoLegKB to retrieve orthology and synteny information**

```
MATCH (g:Gene)-->(o:Orthogroup)<--(g2:Gene)
WHERE g.gene_id = "Vfaba.Hedin2.R1.2g194440"
AND g2.genome in ["vfab", "lcul", "psat", "mtru"]
RETURN g2.gene_id
```

```
MATCH (g:Gene)-->(s:Synteny)<--(g2:Gene)
WHERE g.gene_id = "Vfaba.Hedin2.R1.2g194440"
AND g2.genome in ["vfab", "lcul", "psat", "mtru"]
RETURN g2.gene_id
```

```
MATCH (g:Gene)-->(:Synteny)<--(g2:Gene),
(g)-->(:Orthogroup)<--(g2)
WHERE g.gene_id = "Vfaba.Hedin2.R1.2g194440"
AND g2.genome in ["vfab", "lcul", "psat", "mtru"]
RETURN g2.gene_id
```

**Supplementary File 4. Sequence of *LE* available in Reinecke et al. 2013 and query to identify syntenic blocks between chromosomes Vf05 and Ps05**

```
>U85045.1 Pisum sativum 2-oxoglutarate-dependent dioxygenase mRNA, complete cds
GAATTCACTATGCCTTCACTCTCCGAAGCCTATAGAGCACACCCCGTGCACGTTAACCAC
AAGCACCCTGATTTCAACTCACTTCAAGAACTACCTGAATCTTACAATTGGACTCACCTT
GATGATCACACCCTTATTGATTCCAATAATATTATGAAGGAGAGTACTACTACTGTCCCCG
TTATTGATCTCAATGACCCTAATGCTTCAAAGCTAATAGGACTTGCATGCAAAACATGGG
GGGTGTATCAAGTAATGAACCATGGCATCCCCTTAAGCCTTCTTGAGGATATTCAATGGC
TTGGACAAACACTTTTCTCTCTTCCTTCTCACCAAAAACATAAAGCAACTCGTTCCCCCG
ACGGTGTTTCGGGATATGGCATCGCTCGTATCTCTTCCTTCTTCCCCAAACTCATGTGGTA
TGAGGGATTTACTATCGTCGGATCACCTCTCGACCATTTTCGAGAACTCTGGCCTCAAGA
TTATACCAGATTCTGTGATATTGTCGTGCAATATGATGAAACCATGAAAAAGTTAGCAGG
AACATTAATGTGTCTAATGTTGGACTCTCTTGGTATTACAAAGGAAGATATCAAATGGGC
CGGGTCAAAAGCCCAATTTGAAAAAGCTTGTGCGGCCCTCCAATTAAACTCCTACCCTA
GTTGCCCGGATCCGGATCACGCGATGGGTCTCGCCCCGCACACAGACTCAACATTTTTA
ACCATCCTATCTCAAAACGACATAAGCGGGTTACAGGTTAACCGCGAGGGTTCTGGGTG
GATCACGGTTCCACCGCTCCAAGGAGGTCTGGTCGTCAACGTGGGCGACCTCTTTCATA
TTTTGTCGAACGGGTTATATCCTAGCGTACTCCATCGAGTTTTAGTGAACCGGACCCGTC
AGAGATTTTCCGTTGCCTATTTATATGGCCCCCCTTCCAATGTAGAGATTTGTCCACATGC
AAAATTAATAGGCCCAACAAAACCCCCTCTCTATAGGTCAGTGACATGGAATGAGTACCT
TGGCACAAAAGCAAAACATTTCAACAAAGCACTCTCATCTGTTAGACTTTGTACACCTAT
TAATGGTTTGTTTGATGTAAACGATTCTAACAAAAATAGTGTCCAAGTGGGCTAAATAGG
AATTC
```

```
MATCH (c1:Chromosome)<-[pos1]-(b:Synteny)-[pos2]->(c2:Chromosome)
WHERE c1.chromosome_id = "Vf05"
AND c2.chromosome_id = "Ps05"
RETURN b.block_id, b.nb_genes, c1.chromosome_id, pos1.start, pos1.end, c2.chromosome_id,
pos2.start, pos2.end ORDER BY pos1.start
```

**Supplementary File 5. Cypher query used in OrthoLegKB to retrieve all SEO/SEOR genes and their functional annotation, to confirm that *Vfaba.Hedin2.R1.3g070040* is an ortholog of *MtSEO2***

```
MATCH (g:Gene)-->(r)-->(m:MapMan),
(r)-->(p:Protein)-->(f:FunctionalAnnotation)
WHERE g.genome in ["psat", "vfab", "lcul", "mtru", "cari"]
AND (m.label =~ ".*SEOR.*" OR f.description =~ ".*SIEVE ELEMENT.*")
RETURN DISTINCT g.genome, g.gene_id
```
